# Characterization of Structural Variation in Tibetans Reveals New Evidence of High-altitude Adaptation and Introgression

**DOI:** 10.1101/2020.12.01.401174

**Authors:** Cheng Quan, Yuanfeng Li, Yahui Wang, Jie Ping, Yiming Lu, Gangqiao Zhou

**Affiliations:** Department of Genetics & Integrative Omics, State Key Laboratory of Proteomics, National Center for Protein Sciences, Beijing Institute of Radiation Medicine, Beijing, 100850, P. R. China; Collaborative Innovation Center for Personalized Cancer Medicine, Center for Global Health, School of Public Health, Nanjing Medical University, Nanjing, Jiangsu Province, 211166, P. R. China; Medical College of Guizhou University, Guiyang, Guizhou Province, 550025, P. R. China; Hebei University, Baoding, Hebei Province, 071002, P. R. China

**Keywords:** structural variation, long-read sequencing, Tibetan, high-altitude adaptation

## Abstract

Structural variation (SV) acts as an essential mutational force shaping the evolution and function of the human genome. To investigate the role of SVs in high-altitude adaptation (HAA), we here generated a comprehensive catalog of SVs in a Chinese Tibetan (n = 15) and Han (n = 10) population using the nanopore sequencing technology. Among a total of 38,216 unique SVs in the catalog, 27% were sequence-resolved for the first time. We systemically assessed the distribution of these SVs across repeat sequences and functional genomic regions. Through genotyping in additional 189 genomes, we identified 90 Tibetan-Han stratified SVs and 124 candidate adaptive genes. Besides, we discovered 15 adaptive introgressed SV candidates and provided evidence for a deletion of 335 base pairs at 1p36.32. Overall, our results highlight the important role of SVs in the evolutionary processes of Tibetans’ adaptation to the Qinghai-Tibet Plateau and provide a valuable resource for future HAA studies.

## Introduction

Structural variation (SV) is usually defined as the intra or inter-chromosomal genomic rearrangement acting as a significant mutational force shaping the evolution and function of the genome (Collins et al. 2020). As an essential component of deleterious polymorphisms in humans (Abel et al. 2020), SVs positively impact the formation of population diversity and adaptation (Moan et al. 2019). In the face of high gene flow caused by migration load, SVs can generate incompatible alleles allowing the beneficial variant to reduce the risk of recombination with maladapted genomic backgrounds, thereby maintaining allele frequency (AF) clines towards environmental gradients (Moan et al. 2019; Wellenreuther et al. 2019).

The adaptation of Tibetan highlanders to the Qinghai-Tibet Plateau with an average elevation of over 4,000 meters is a representative case of anatomically modern human (AMH) conquering new environmental conditions (Huerta-Sánchez et al. 2014). Previous studies usually used single nucleotide variants (SNVs) to search for selection evidence in the Tibetan genome and successfully identified two essential genes involved in the HIF-1 (hypoxia-inducible transcription factor 1) pathway, *EPAS1* (endothelial PAS domain protein 1) and *EGLN1* (egl-9 family hypoxia inducible factor 1) (Xiang et al. 2013; Huerta-Sánchez et al. 2014; Deng et al. 2019). The adaptive alleles of these two genes could help maintain the hemoglobin concentration so that red cells would not be overproduced at high altitude (Hu et al. 2017; Yang et al. 2017). However, few studies have examined the role of SVs underlying high-altitude adaptation (HAA). In an earlier study (Lou et al. 2015), a 3.4 kilobase (kb) Tibetan-enriched deletion (TED) located at 80 kb downstream of *EPAS1* had been identified using microarray data, indicating that SVs also play a role in the process of Tibetans’ adaptation to the plateau. Besides, the discovery of a particular Tibetan-Denisovan haplotype motif characterized by derived alleles from five SNVs in the *EPAS1* region indicated shared genetic drift between Tibetans and extinct non-AMH populations (Yi et al. 2010; Huerta-Sánchez et al. 2014; Lu et al. 2016). Although several recent studies have found that SVs originated from archaic hominins and introgressed into AMH substantially contribute to local population adaptation (Hsieh et al. 2019; Almarri et al. 2020), little is known of adaptive introgressed SVs in Tibetans so far.

In recent years, long-read sequencing (LRS) technologies, such as PacBio single-molecule real-time (SMRT) sequencing and nanopore sequencing, have been introduced to resolve SVs in the human genome (Stancu et al. 2017; Audano et al. 2019). Long and contiguous reads generated from these emerging technologies can span SVs’ breakpoints, enabling more accurate alignment to repetitive sequences (Coster et al. 2019). Thus, LRS makes it possible to detect previously unresolved SVs with high sensitivity (Sedlazeck et al. 2018a; Chaisson et al. 2019; Ho et al. 2020), facilitating a better evaluation of SVs’ impact on HAA. A recent study sequenced one single adult male of native Tibetan ancestry using the PacBio SMRT sequencing platform, and identified a deletion of 163 base pairs (bp) located in an intronic region of *MRTFA* (myocardin related transcription factor A), which is associated with lower systolic pulmonary arterial pressure in Tibetans (Ouzhuluobu et al. 2019). However, it is still necessary to use population-based detection methods to explore SVs related to HAA.

In this study, we sought to comprehensively characterize the complete spectrum of SVs in a Chinese Tibetan and Han population using the nanopore sequencing technology. We performed a population-based SV detection in order to include as many common variants as possible. Among a total of 38,216 unique SVs in the catalog, we identified 90 Tibetan-Han stratified SVs and 124 candidate adaptive genes, providing valuable resources for future HAA studies. Furthermore, we scanned signatures of natural selection and archaic introgression around the population-stratified SVs to explore how SVs introgressed from hominins could facilitate HAA.

## Results

### Discovery of SVs in a Tibetan and Han population by long-read sequencing

Long-read sequencing data were generated from 15 Tibetan and 10 Han genomes using the nanopore sequencing (Supplemental Table S1). For each genome, we detected SVs relative to GRCh37 human reference assembly using a multi-platform pipeline (Methods). On average, 15,813 SVs per sample were identified and were then merged into a non-redundant set of 41,792 SVs (Fig. 1A and 1B). Consistent with a previous study (Audano et al. 2019), the size of the non-redundant set grew rapidly at the beginning and gradually slowed down with the increase of samples, suggesting that a considerable proportion of common variations were detected in this Chinese Tibetan and Han population. Then, we filtered out 3,576 unreliable genotypes with a hard threshold and obtained a total of 38,216 unique SVs (Methods). Nearly 74% (n = 27,994) of these SVs have breakpoints within an interval of 100 bp across different samples (Supplemental Fig. S1A). All subsequent analyses were then performed on this final set of SVs, which comprise 19,441 deletions, 17,424 insertions, 721 duplications, 442 inversions, and 188 translocations (Fig. 1C; Supplemental Table S3). Contrary to the results observed using PacBio SMRT-SV (Audano et al. 2019), we identified more deletions than insertions using nanopore sequencing. This phenomenon might result from base-calling errors of homopolymer regions, which is still the main drawback of nanopore sequencing (Stancu et al. 2017; Jain et al. 2018; Sedlazeck et al. 2018b; Coster and Broeckhoven 2019). Meanwhile, we identified a relatively small number of duplications, mainly because the majority of common duplications should be classified as multi-allelic SVs (Sudmant et al. 2015), which is beyond the scope of this study.

**Fig. 1.**
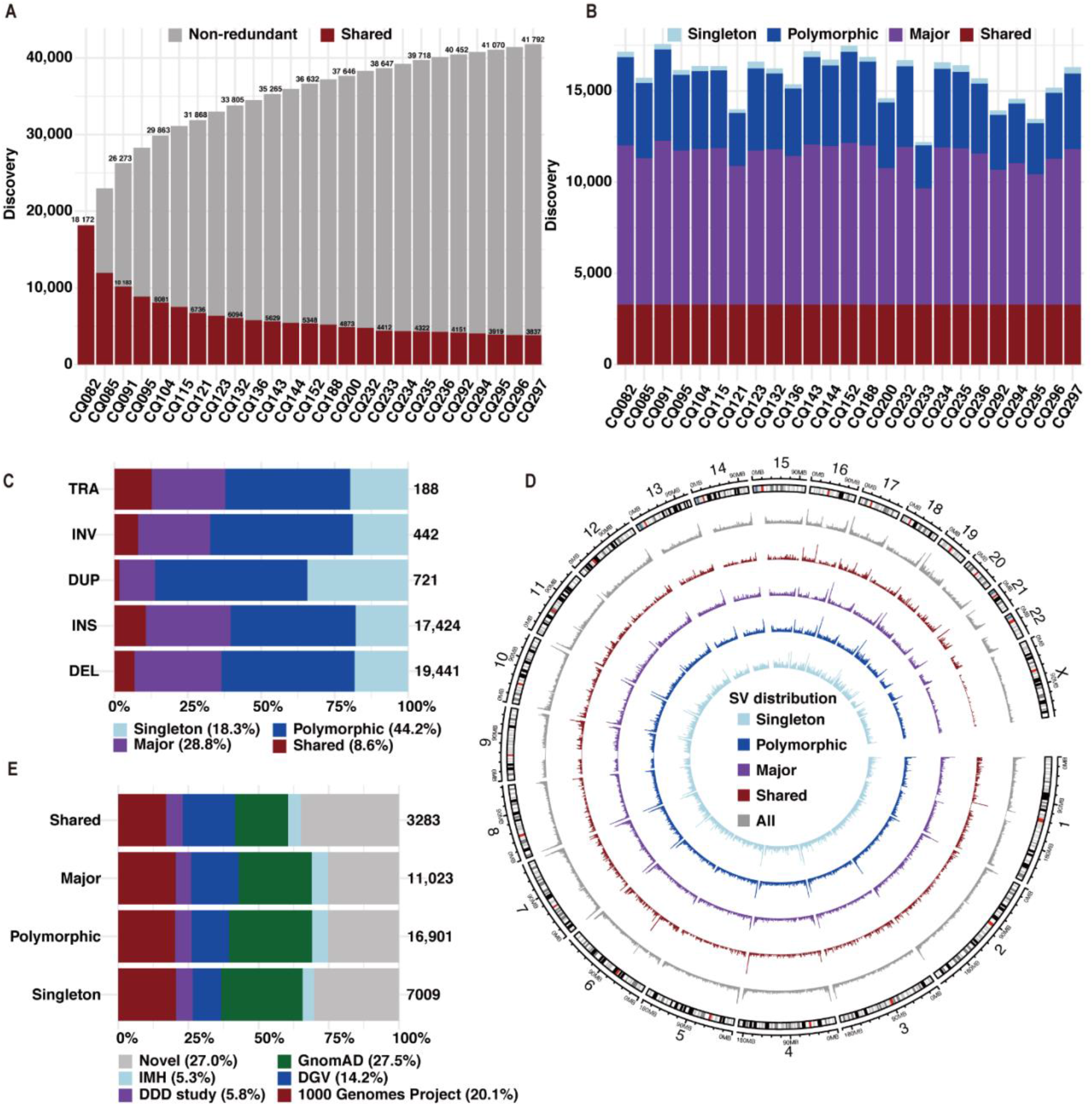
Discovery of structural variations in 15 Tibetan and 10 Han samples. (A) Structural variations (SVs) discovered from each sample were merged into a non-redundant set. Shared SVs are shown as red portions of each bar. (B) The number of SVs for each discovery category is shown per sample, including shared (identified in all samples), major (identified in ≥ 50% of samples), polymorphic (identified in > 1 sample), and singleton (identified in only one sample) SVs. (C) The frequencies for each SV type: translocation (TRA), inversion (INV), duplication (DUP), insertion (INS), and deletion (DEL). (D) The circular layout of SV distribution for each discovery category. (E) Proportions for SVs discovered in previously published SV calls for each discovery category. IMH represents a common disease trait mapping study published by Ira M. Hall’s lab. DDD study represents the Deciphering Developmental Disorders Study.

As suggested in a previous report (Audano et al. 2019), we classified these SVs into four categories: shared (identified in all samples), major (identified in ≥ 50% of samples), polymorphic (identified in > 1 sample), and singleton (identified in only one sample) SVs. Consistent with the results observed using PacBio SMRT-SV (Audano et al. 2019), nearly 40% (n = 14,306) of the SVs were identified as major or shared variants, and the proportion of shared insertions was higher than that of deletions (Fig. 1C). The frequencies of SVs in all categories decreased markedly with length, and > 88% (n = 33,680) of these SVs have a length ranging from 50 bp to 1 kb (Supplemental Fig. S1B and S1C). As reported in previous studies (Audano et al. 2019; Beyter et al. 2019), we estimated a fourfold increase in SV density within the last five megabase (Mb) of chromosome arms (p-value = 0.001), with the largest increase for major variants (fold change [FC] = 5.48) and the lowest for shared variants (FC = 4.18) (Fig. 1D and Supplemental Fig. S2).

We compared our non-redundant set with previously published SV calls identified from thousands of genomes and calculated the maximum variant frequency in these datasets (Methods). We found that 27.0% (n = 10,308) of the SVs discovered in our data are novel ones that have not been sequence-resolved before (Fig. 1E). Moreover, there were 842 shared variants and 2,310 major variants defined as rare mutations (maximum AF < 0.01) in previous studies (Supplemental Fig. S1D), suggesting an increased sensitivity for SV detection using the nanopore sequencing technology (Sedlazeck et al. 2018a; Chaisson et al. 2019; Ho et al. 2020).

### Distribution of SVs across repeat sequences and functional genomic regions

We found that nearly 80% of the breakpoints of SVs overlapped with repetitive elements, of which most are interspersed repeats (56.8%) and tandem repeats (21.3%) (Fig. 2A). Compared to the random background, we found significant positive enrichment of short interspersed nuclear elements (SINEs) (FC = 1.22, p-value = 0.009) and clear depletion of other interspersed repeats (Fig. 2B), such as long interspersed nuclear elements (LINEs) (FC = 0.78, p-value < 0.001) and long terminal repeats (LTRs) (FC = 0.80, p-value < 0.001). These results indicated that the relatively small size of SINEs makes them more neutral than other interspersed repeats, so they manage to act as an important source of genetic diversity associated with the formation of SVs (Zhao et al. 2016; Payer et al. 2017). Consistent with previous studies (Beyter et al. 2019; Coster et al. 2019), the SV length distribution profiles showed noticeable peaks at 300 bp, corresponding to SINEs, as well as 6 kb, corresponding to LINEs (Fig. 2A). To our surprise, we also found a considerable number of SV events around 300 bp overlapped with LINEs, especially for insertions (Supplemental Fig. S3A and S3B). Previous studies have reported that the Alu element is a non-autonomous retrotransposon belong to SINE and has a length of ~300 bp. Therefore, the Alu element acquires trans-acting factors from LINE-1 (Deininger 2011), which could result in overlaps of SINEs and LINEs in modern human genomes.

**Fig. 2.**
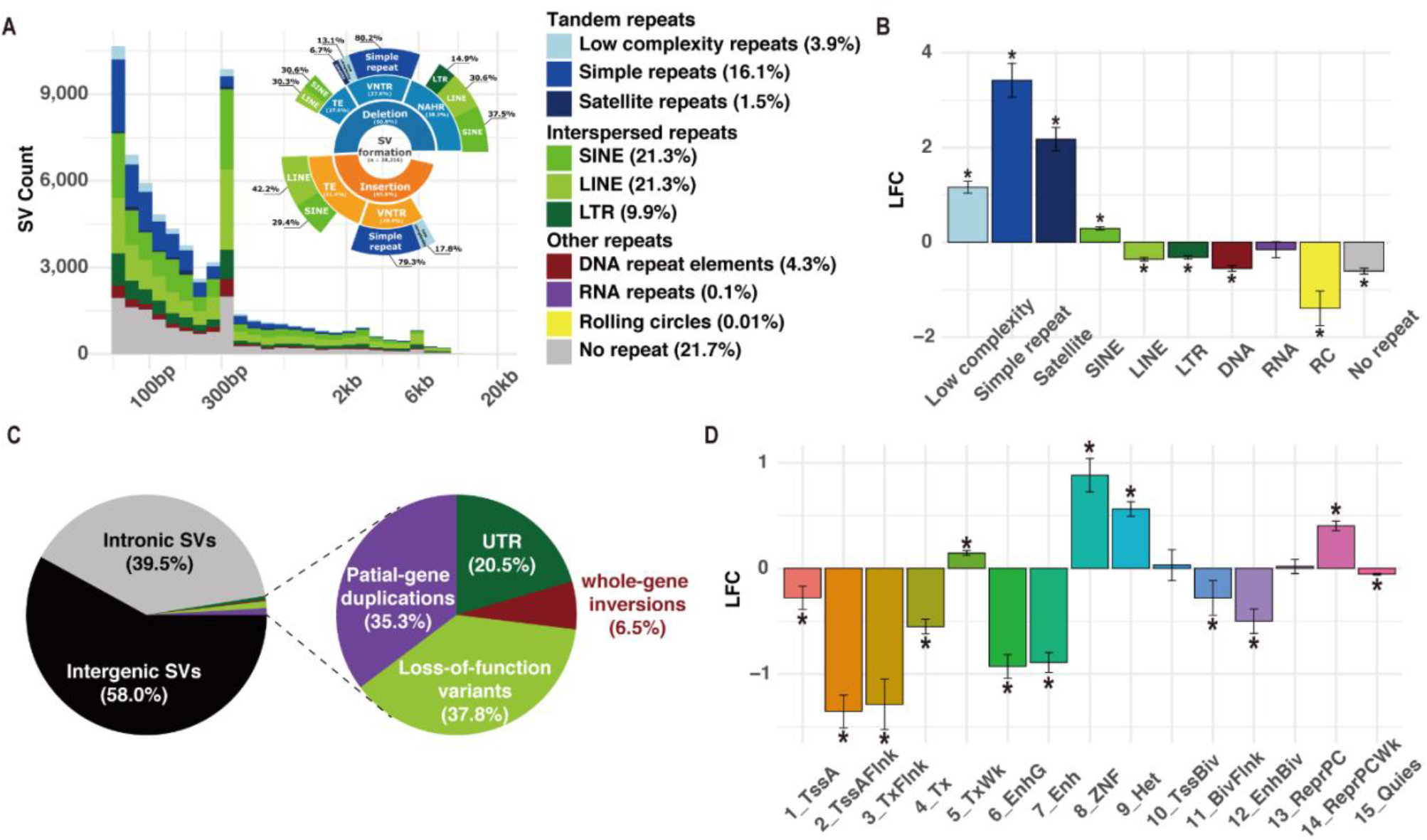
Distribution of structural variations across repeat sequences and functional genomic regions. (A) Distribution of insertions and deletions classified by intersected repeat elements. The formation mechanism was classified into three categories, including nonallelic homologous recombination (NAHR), variable number tandem repeats (VNTR), and transposable elements (TE). (B) Log2 fold change (LFC) of enrichment analysis for repeat elements intersected with breakpoint junction sequences. ^*^denotes significant enrichment with empirical p-value < 0.05. (C) Proportions for potential effects on coding sequences. (D) LFC of enrichment analysis for structural variations intersected with 15-core chromatin states from RoadMap.

Then, we used a simplified pipeline to infer the SV formation mechanisms according to the profile of breakpoint junction sequences (Methods; Fig. 2A). Nonallelic homologous recombination (NAHR) was the dominated formation mechanism (38.3%, n = 7,433) among all deletion events (< 1 Mb, n = 19,416), consistent with previous observations of SV hotspots (Mills et al. 2011). Further classification of deletions showed that 37.5% (n = 2,787) of the NAHR events overlapped with SINE on both sides, again suggesting that SINE plays an essential role in the formation of SVs. Variable number tandem repeats (VNTRs) were another contributor to SV formation, accounting for 27.6% (n = 5,350) of deletions and 30.4% (n = 5,296) of insertions (< 1 Mb, n = 17,424). Additionally, 41.4% (n = 7,221) of insertions were mediated by transposable elements (TE), and approximately 71.7% (n = 5,177) out of them were annotated as LINE or SINE-mediated ones.

As SVs usually have deleterious effects when occurring in functional sequences (Hurles et al. 2008; Lin and Gokcumen 2019), we systemically evaluated possible functional impacts of major and shared SVs on both the coding and non-coding regions. First, we found that most SVs are located in introns or intergenic regions (Fig. 2C). Compared to the genomic background (Supplemental Table S4), SVs which were predicted to cause genic loss of function (LoF) showed a significant depletion (p-value < 0.001), while the intergenic SVs showed significant enrichment (p-value = 0.009). Further analysis of the nearest transcription start site (TSS) revealed that SVs rarely locate near the genes that are intolerant to either LoF (p-value < 0.001) or missense mutations (p-value = 0.049). Besides, we found significant depletion of SVs overlapped with binding clusters of transcriptional repressor CTCF (p-value < 0.001) and boundaries of topologically associating domain (TAD) (p-value < 0.001). In addition, SVs are also strongly depleted in active chromatin regions and enriched in inactive regions (Fig. 2D). An exception is that a significant enrichment of SVs was found in weakly transcripted regions (TxWK, p-value = 0.009; Fig. 2D).

### Population structure and Demographic inference

To further study the population distribution of SVs discovered by LRS, we aggregated a human genome diversity panel constructed from 189 genomes using next-generation sequencing (NGS) data, including 81 Tibetans, 84 Hans, and 24 Biaka samples in the Central African Republic (Methods; Supplemental Table S2). Targeted genotyping of SVs using NGS data is still a nontrivial challenging task (Sedlazeck et al. 2018a; Chen et al. 2019). In this study, we applied a graph-based approach (Chen et al. 2019), and 83.8% (31,881/38,028) of the SVs were successfully genotyped in at least 95% of samples (Fig. 3A), which means the missing rate (MR) of genotyping is less than 0.05. However, only 53.2% (16,953/31,881) of these successfully genotyped SVs were supported by at least one sample (Fig. 3A), indicating that nearly half of the SVs discovered by LRS can not be recalled by NGS (genotyped AF = 0). Although the graph-based method reduced the MR of genotyping, it still resulted in an unsatisfying recall rate, similar to previous studies (Audano et al. 2019; Beyter et al. 2019). Further analysis showed that 28.6% (4,263/14,928) of these SVs, which failed to be recalled by NGS, were classified as major or shared SVs in the original LRS calls (Fig. 3A). This result indicated that the low recall rates are unlikely to be solely caused by rare mutations or somatic artifacts, at least for these common SVs discovered by LRS. Besides, 56.7% (8,465/14,928) of these SVs overlapped with repeated sequences or segmental duplications, and another 19.5% (2,917/14,928) has repetitive breakpoint junction sequences (Supplemental Fig. S4A). These results suggested that the relatively low resolution in the repeated regions may be a significant flaw in SV genotyping with NGS data.

**Fig. 3.**
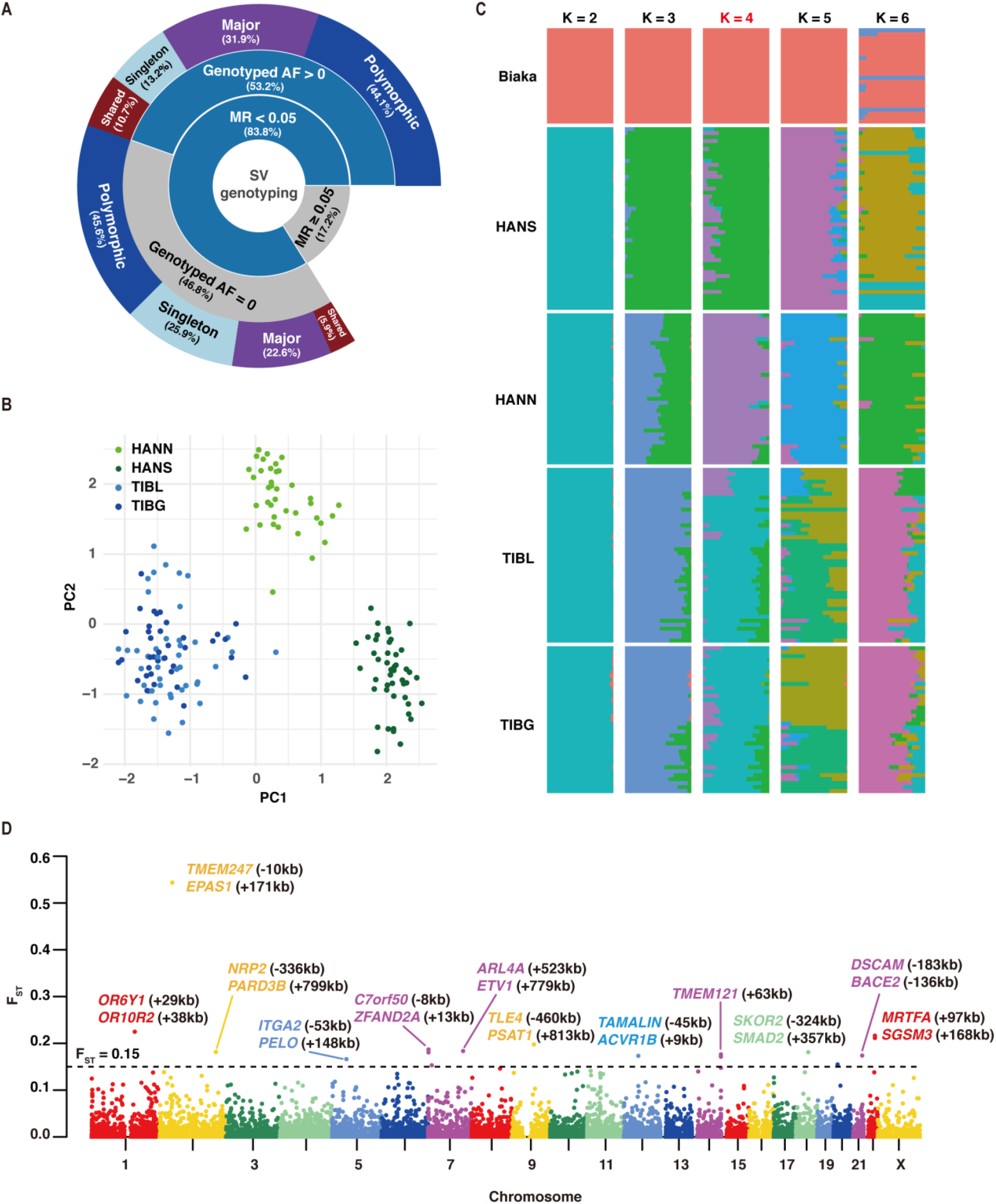
Population structure analysis using short-read data. (A) Genotyping results for short-read sequencing data. Discovery categories for original LRS calls were shown for structural variations (SV) with genotyped missing rate (MR) < 0.05, including shared (identified in all samples), major (identified in ≥ 50% of samples), polymorphic (identified in > 1 sample), and singleton (identified in only one sample) SVs. (B) Principle component analysis (PCA) of SV genotypes for Hans in North China (HANN) or South China (HANS), and Tibetans living above (TIBG) or below (TIBL) 4,000 meters. (C) Admixture analysis of Tibetans and Hans, using the Biaka populations as an outgroup. The minimum cross-validation error occurs when K = 4. (D) Manhattan plot for the window-based fixation index (F_ST_) statistics (Tibetans *vs.* Hans). Linear nearest genes to top candidate regions (F_ST_ > 0.15) are marked in the corresponding colors. Distances from the middle of the regions to the transcription start site of the genes are listed in brackets.

Taking advantage of these successfully recalled SVs (n = 16,953, AF > 0 and MR < 0.05), we further investigated the population structure. First, we artificially divided Chinese samples into four subgroups based on geography, including Hans in North China (HANN) or South China (HANS), and Tibetans living above (TIBG) or below (TIBL) 4,000 meters. Meanwhile, Biaka samples were introduced as an outgroup (Supplemental Table S2). While large proportions of rare SVs were specific to individual groups, almost all SVs were shared across these groups at AF > 2.5% (Supplemental Fig. S4B). We found little difference in SV-homozygosity across populations (Supplemental Fig. S4C). HANS and Biaka populations exhibited slightly less homozygous than the other groups. Biaka samples exhibited 10% less heterozygous from Chinese populations (mean of 3,203 *vs.* 3,577, p-value < 0.001), indicating that many genotyped SV events were unique to Chinese populations. We did not find any significant differences between TIBG and TIBL, while the heterozygosity of HANS was significantly higher than HANN (mean of 3,644 *vs.* 3,522, p-value < 0.001). Principle component analysis (PCA) also revealed that Biaka and Chinese samples were clustered in separated groups (Supplemental Fig. S4D). Restricting the analysis to Chinese populations unveiled three major clusters (Fig. 3B), with the first PCA axis separating Tibetans from Hans, and the second axis separating HANS from HANN. The best-fitting admixture model indicated by the minimum cross-validation error was K = 4 (Fig. 3C), depicting four primary ancestral components, including Biaka, TIB, HANN, and HANS.

After the major population structure (TIB-HANN-HANS) was confirmed, we used whole-genome SNVs to estimate the demographic history of the three populations (Methods). We tested three models with symmetric migrations to describe simplified evolution paths (Supplemental Table S5). After rounds of optimization, the second model with no size changes yielded the highest likelihood, describing a more recent divergence between HANN and HANS than between HAN and TIB. Specifically, the best-fit demographic model (Supplemental Fig. S5A and S5B; Supplemental Table S7) indicated that the ancestors of Tibetans and Hans remained isolated until ~17,000 years ago (95% confidence interval [CI]: 14,776 to 20,502 years), while HANN and HANS diverged much more recently, ~770 years ago (95% CI: 727 to 810 years). These results confirmed the recent migration to the Qinghai-Tibet Plateau after the intense cold period during the Last Glacial Maximum (LGM, ~19,000 to 25,600 years before present) (Lu et al. 2016). If we ignored the internal genetic divergence in Han populations and included Yoruba samples in Ibadan (YRI) to construct another population model YRI-TIB-HAN (Methods; Supplemental Table S6), the best-fit demographic model (Supplemental Fig. S5C and S5D; Supplemental Table S7) indicated that the ancestors of Africans and Chinese diverged ~88,000 years ago (95% CI: 82,872 to 94,398 years), followed by TIB-HAN divergence ~22,000 years ago (95% CI: 19,763 to 26,184 years). In this case, it takes a longer time for neutral variants to reach the level of variation within the Han populations. As a result, the divergence between Tibetans and Hans is much closer to the first major prehistoric migration into the plateau in the Upper Paleolithic (~30,000 years before present), supported by both archaeological and genetic studies (Aldenderfer 2011; Qi et al. 2013).

### Population-stratified SVs and candidate adaptive genes

In order to discover candidate adaptive variants, we calculated the fixation index (F_ST_) between Tibetans and Hans using the NGS data (Methods). A total of 90 Tibetan-Han stratified SVs (F_ST_ > 0.1) were identified, including 75 deletions and 15 insertions (Fig. 3D; Table 1 and Supplemental Table S9). Less than one-fifth of these SVs (15/90) have a length > 1 kb, and the largest event is a ~61 kb deletion at chromosome 20p11. Almost all the population-stratified SVs were located in introns or intergenic regions, except for three deletions leading to the ablation of pseudogene transcripts (Supplemental Table S9). Consistent with previous reports (Lou et al. 2015; Ouzhuluobu et al. 2019), the top candidate variant with the maximum F_ST_ is the 3.4 kb TED located in the intergenic region between *EPAS1* and *TMEM247* (transmembrane protein 247). We also confirmed the 163 bp Tibetan-Han stratified deletion in the intronic region of *MRTFA* (Ouzhuluobu et al. 2019).

**Table 1.** Summary of top 13 population-stratified structural variations (SVs) with F_ST_ > 0.15

| No. | SV Location (GRCh37) | Type | Length<br>(bp) | $F_{ST}$ | Genotyped AF | | | Candidate Affected<br>Genes |
| --- | --- | --- | --- | --- | --- | --- | --- | --- |
|  |  |  |  |  | Tibetan | Han | Biaka |  |
| 1 | chr2:46,694,272-46,697,680 | Deletion | 3,408 | 0.54 | 0.58 | 0.02 | 0.00 | *TMEM247* and other 8 genes |
| 2 | chr6:57,392,157-57,392,158 | Insertion | 4,197 | 0.24 | 0.61 | 0.24 | 0.63 | *PRIM2* |
| 3 | chr1:158,488,405-158,488,672 | Deletion | 267 | 0.22 | 0.84 | 0.51 | 0.56 | *OR10Z1* |
| 4 | chr22:40,935,457-40,935,620 | Deletion | 163 | 0.22 | 0.51 | 0.17 | 0.00 | *MRTFA*, *EP300* |
| 5 | chr9:81,725,770-81,726,090 | Deletion | 320 | 0.20 | 0.63 | 0.30 | 0.56 | *TLE4* |
| 6 | chr7:1,185,068-1,187,658 | Deletion | 2,590 | 0.19 | 0.77 | 0.45 | 0.92 | *ZFAND2A*, *GPER1* |
| 7 | chr7:127,766,419-127,766,732 | Deletion | 313 | 0.18 | 0.62 | 0.30 | 1.00 | *LRRC4* |
| 8 | chr2:206,210,503-206,210,504 | Insertion | 3,345 | 0.18 | 0.73 | 0.41 | 0.13 | *PARD3B* |
| 9 | chr18:45,099,900-45,100,212 | Deletion | 312 | 0.18 | 0.24 | 0.02 | 0.48 | *SKOR2* |
| 10 | chr14:106,055,915-106,056,047 | Deletion | 132 | 0.18 | 0.10 | 0.36 | 0.00 | *TMEM121*, *IGHA2* |
| 11 | chr21:42,402,890-42,403,212 | Deletion | 322 | 0.17 | 0.70 | 0.39 | 1.00 | *BACE2* |
| 12 | chr12:52,354,758-52,354,891 | Deletion | 133 | 0.17 | 0.60 | 0.29 | 0.04 | *ACVR1B* |
| 13 | chr5:52,231,784-52,232,117 | Deletion | 333 | 0.17 | 0.63 | 0.89 | 0.90 | *ITGA1* |

$F_{ST}$ , fixation index between Tibetans and Hans; AF, allele frequency; bp, base pairs.

SVs could alter the transcription of genes by affecting regulatory elements in non-coding regions (Fudenberg and Pollard 2019; Jakubosky et al. 2020; Shanta et al. 2020), which is called position effects (Spielmann et al. 2018). When occurring near genes, SVs exhibit remarkable regulatory effects and are more likely to act as expression quantitative trait locus (eQTL) than SNVs in distal regions (Jakubosky et al. 2020). In order to identify all candidate adaptive genes that tend to be affected by the Tibetan-Han stratified SVs, we considered three types of evidence-supported genes, including linear overlapping/nearest genes, three-dimensional (3D) affected genes, and linkage disequilibrium (LD)-linked eQTL genes (eGenes). Finally, we discovered a total of 124 protein-coding genes (Fig. 4A and 5D; Supplemental Table S10). Among these potentially affected genes, 36 genes have been reported in previous studies to be relevant to HAA, and another 64 genes have traits directly or indirectly related to hypoxia (Supplemental Table S10). Here, we briefly summarize the potential functional effects of some representative SVs on these candidate adaptive genes.

**Fig. 4.**
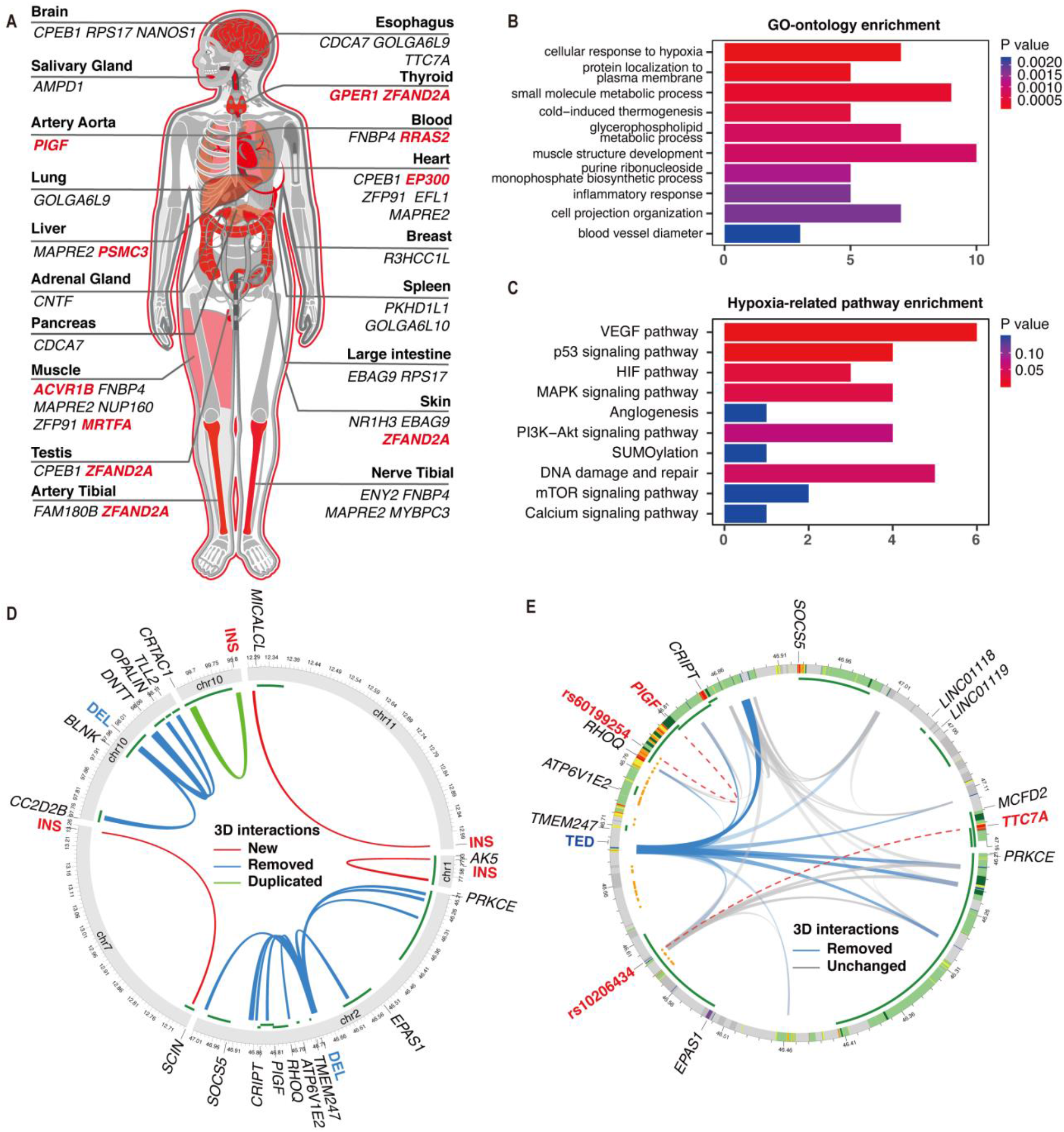
Candidate adaptive genes. (A) All linkage disequilibrium (LD)-linked genes across tissues. Known adaptive genes were marked in red color. (B) Gene-Ontology (GO) and (C) Hypoxia-related pathway enrichment analysis for 124 candidate genes. (D) All three-dimensional (3D) interactions and the corresponding genes affected by 90 population-stratified structural variations (SVs) (fixation index [F_ST_] > 0.1 between Tibetans and Hans). The outermost green layer represents the transcription regions of affected genes. Deletions (DEL) will remove overlapped interactions (blue), and insertions (INS) could create a new anchor (red) or duplicate nearby anchors and then construct new interactions with the closest anchors (green). (E) Profile of all CTCF interactions near the 3.4 kilobase (kb) Tibetan-enriched deletion (TED). The outermost layer shows the 15-core chromatin states from RoadMap. The inner green layer represents the transcription regions of affected genes. Among single nucleotide variants (SNVs) highly linked with TED (orange points), rs60199254 and rs10206434 act as expression quantitative trait locus (eQTL) exhibiting relationships with expression levels of *PIGF* and *TTC7A,* respectively (red dashed lines).

We first assessed the functional roles of the two known adaptive SVs. We found that the 3.4 kb TED, which has been reported to affect the function of *EPAS1* (Lou et al. 2015), is able to modify the high-order chromatin structure by removing a nearby chromatin loop anchor that mediated interactions between regulatory elements and multiple distant genes (Fig. 4E). For example, the TED could remove a convergent interaction between 46.68-46.88 Mb in chromosome 2, which will disable the interactions between many strong enhancers/active promotors and five continuous protein-coding genes within this region (Fig. 4E), including *TMEM247*, *ATP6V1E2*, *RHOQ*, *PIGF*, and *CRIPT*. Through the discovery of Tibetan-Han stratified SNVs in this region, it has been proved that all these candidate genes were relevant to HAA (Huerta-Sánchez et al. 2014; Deng et al. 2019). Furthermore, LD-linked eQTL tests also confirmed the relationship between the TED and *PIGF* (Fig. 4E and Supplemental Fig. S6A; Supplemental Tables S11 and S12). *PIGF* (phosphatidylinositol glycan anchor biosynthesis class F) is an essential member of the *VEGF* (vascular endothelial growth factor) family and facilitates angiogenesis in Tibetans (Deng et al. 2019). Therefore, the TED is likely to regulate many adaptive genes besides *EPAS1*. Similarly, we found that the 163 bp deletion in the intronic region of *MRTFA* was also involved in a high-confidence LD-linked eQTL relationship with *EP300* (E1A binding protein p300) (Supplemental Fig. S6B; Supplemental Tables S11 and S12), which encodes a co-activator of *HIF1A* and can stimulate other hypoxia-induced genes (Deng et al. 2019). Together, these results provide new insights into the two known SVs and suggest population-stratified SVs could affect multiple adaptive genes far from their breakpoints.

Besides, we found that a substantial number of adaptive genes (n = 36) identified by SNVs are potentially influenced by novel SVs. For example, a 2,590 bp deletion at 7p22.3 (chr7:1,185,070-1,187,063) is located nearby a region with reported selection signatures (Deng et al. 2019). Consistent with previous reports (Yang et al. 2017; Deng et al. 2019), four SNVs (rs7805591, rs2949174, rs2140578, rs2949172) highly linked to this deletion (LD > 0.8) showed significantly different frequencies between Tibetans and Hans (all F_ST_ > 0.16; Supplemental Table S13). These adaptive SNVs were identified as eQTLs exhibiting negative associations with the expression levels of *ZFAND2A* (zinc finger AN1-type containing 2A) and *GPER1* (G protein-coupled estrogen receptor 1) (Fig. 4A and Supplemental Fig. S6C; Supplemental Tables S11 and S12). Both *ZFAND2A* and *GPER1* locate close to this deletion and are associated with HIF1A. *ZFAND2A* is predicted to be a target gene of the HIF1A transcription factor using known transcription factor binding site motifs from the TRANSFAC database (Matys et al. 2006; Rouillard et al. 2016). Moreover, *ZFAND2A* was recently reported to undergo positive selection and exhibit significantly down-regulated expression in the Tibetan pig (Wu et al. 2019). Meanwhile, activation of *GPER1* by 17β-estradiol (E2) and the specific agonist G-1 will trigger a GPER1-EGFR-MAPK1-FOS signaling pathway, leading to the increased expression level of VEGF through upregulation of HIF1A (Francesco et al. 2014; Xiang et al. 2018). These results suggested that the regulatory mechanism of adaptive genes is complex and could be affected by both SNV and SV.

Lastly, we performed functional enrichment to select novel adaptive genes for HAA. Gene-Ontology enrichment analysis (Fig. 4B; Supplemental Table S14) showed that these candidate genes were significantly involved in the cellular response to hypoxia, cold-induced thermogenesis, blood vessel diameter regulation, and etc. Besides, enrichment analysis of hypoxia-related pathways indicated that candidate genes are most significantly enriched in the VEGF pathway (Fig. 4C; Supplemental Table S15), which could stimulate the proliferation of endothelial cells (ECs) and promote angiogenesis (Senger et al. 2002). Among the novel candidates responding to hypoxia, *CPEB1* (cytoplasmic polyadenylation element binding protein 1) encodes a protein that binds to the 3′-UTR of *HIF1A* mRNA and affects its translational efficiency; when activated by hypoxia and insulin, CPEB1 will increase the protein expression of HIF1A (Hägele et al. 2009). Another candidate, *ITGA1* (integrin subunit alpha 1), encodes a positive regulator of blood vessel diameter; ITGA1 usually dimerizes with ITGB1 and forms a cell-surface receptor, which facilitates attachment of dermal microvascular ECs and collaborates with VEGF in promoting MAPK activation and angiogenesis (Senger et al. 2002). These results demonstrated that a considerable number of genes affected by population-stratified SVs are involved in the biological functions contributing to HAA, suggesting they are novel adaptive genes.

### Signatures of positive selection and archaic introgression

As described in previous studies (Hsieh et al. 2019; Almarri et al. 2020), the population-stratified SVs can originate from de-novo mutations or standing variations that have been introgressed from other hominins and were then subject to natural selection or demographic processes. To distinguish these two hypotheses, we searched for signatures of selection and archaic hominin introgression using population genetic statistics towards SNVs from the sequences surrounding the Tibetan-Han stratified SVs (Methods). Neutral variations from parametric coalescent simulations based on the best-fit demographic model for Tibetans (Supplemental Fig. S5 and S7; Supplementaty Table S8) were considered as the null expectation to estimate the significance of selection and introgression at individual loci (Methods). Using evaluation methods described in previous studies (Hsieh et al. 2016; Hsieh et al. 2019), we found that our whole-genome coalescent simulations could recapitulate the mutation patterns of SNVs and exhibited the ability to identify non-neutral variants (Methods; Supplemental Fig. S8 and S9). We determined the selection and introgression candidates by checking whether they are close (< 500 kb) to a significant signature. Among the 90 Tibetan-Han stratified SVs, we identified signatures of natural selection around 20 SV loci (p < 0.05, integrated haplotype homozygosity score [iHS]) and archaic introgression around 61 loci (p < 0.05 for both D-statistic and f_d_-statistic), including 52 and 39 loci using Neanderthal and Denisovan genomes as the archaic reference respectively. Three-quarters of the selection candidates (15/20) also have introgression signals at the flanking sequences, which were marked as adaptive introgressed SVs (Fig. 5; Supplemental Tables S9 and S16).

**Fig. 5.**
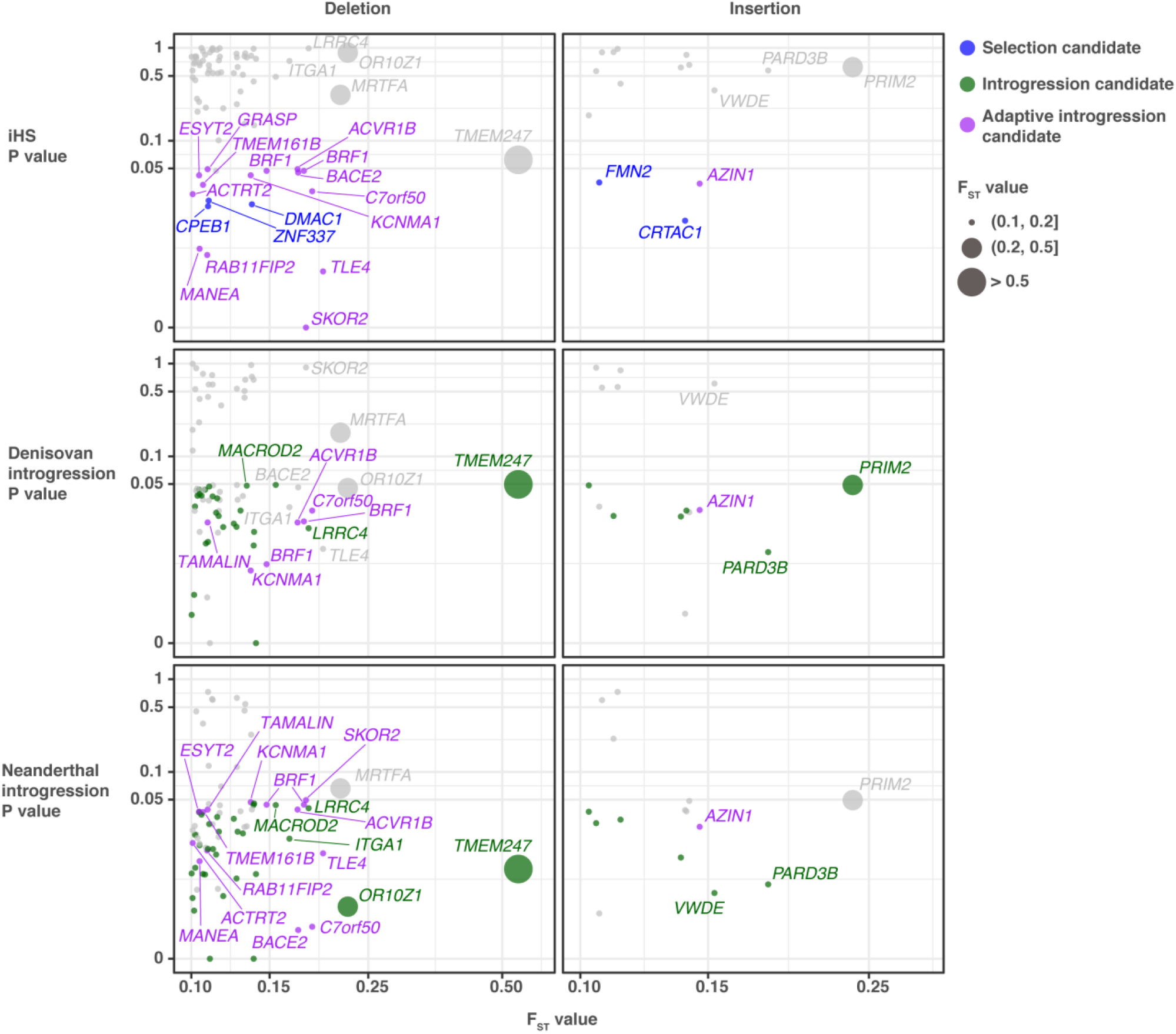
Candidate adaptive and archaic introgressed structural variations. Joint distributions of fixation index (F_ST_) statistics for the population-stratified structural variations (SVs) (x-axis), p-values for positive selection (integrated haplotype homozygosity score [iHS], top row), Denisovan introgression (f_d_-statistic, middle row), and Neanderthal introgression (f_d_-statistic, bottom row). SVs that show signatures of both selection and introgression (purple circles) are distinguished from the variants that show signatures of selection (blue circles) or introgression (green circles) only. The p-values point to the most significant and closest windows to SVs (±500 kb).

To further validate these SVs in ancient genomes, we applied the same graph-based genotyping approach to archaic hominins and chimpanzees (Methods). Among 15 adaptive introgression candidates, three deletions were supported by at least one ancient genome with short-read sequencing data (Supplemental Table 16), including a 335 bp deletion at 1p36.32 (chr1:2,919,030-2,919,365), a 2,590 bp deletion at 7p22.3 (chr7:1,185,070-1,187,063), and a 322 bp deletion at 21q22.2 (chr21:42,402,890-42,403,212). Among these three candidates, the deletions at 7p22.3 and 21q22.2 caused copy number losses in both Biaka and ancient populations (Supplemental Fig. S10A and S11A). Meanwhile, both deletions exhibited high genotyped variant frequencies in the Biaka population (AF = 0.92 and 1, respectively; Supplemental Fig. S10B and S11B). Therefore, it is difficult to determine whether these two deletions are derived from ancient humans, because Neandertals and Dennisovas should not have obvious genetic communications with African ancestors (Hsieh et al. 2019; Almarri et al. 2020).

The deletion at 1p36.32 (chr1:2,919,030-2,919,365) was found only in the Chinese and ancient genomes (Fig. 6B). No evident copy number losses around this deletion were found in either Biaka genomes or Chimpanzees (Fig. 6A), suggesting a high-confidence introgressed SV locus. Meanwhile, the flanking SNVs exhibited significant positive cross-population Extended Haplotype Homozygosity (XP-EHH), suggesting that this region might undergo positive selection in the Tibetans (Fig. 6C; Supplemental Tables S17 and S18). It is worth noting that this SV occurred less frequently in Tibetans (AF = 0.06) than in Hans (AF = 0.23) (Fig. 6B), suggesting that it is the ancestral allele subject to positive selection. Although the allele frequency was relatively lower than other Tibetan-Han stratified SVs, this deletion exhibited high rates of heterozygosity in both Tibetans (100%) and Hans (88%), indicating a considerable variant frequency in Chinese populations (Fig. 6B). We further found that this SV exists in populations from the 1000 Genomes Project and has the highest variant frequency in East Asians (AF = 0.27) and the lowest in Africans (AF = 0.06). Therefore, we extracted 90 flanking SNVs of this deletion with positive selection signals (XP-EHH > 2) and detected the differences between haplotypes of the Tibetan, Han, and African populations. Through hierarchical clustering and network analysis (Fig. 6D and 6E), we found that all haplotypes with the deletion in Tibetans and Hans were clustered together, as well as the haplotypes with or without the deletion in the ancient genomes. In contrast, all haplotypes with the deletion in YRI were clustered with other haplotypes without the deletion, indicating a distinct pattern from the Chinese population. As expected, the beneficial standing ancestral allele co-exists with complex genomic backgrounds, which resulted in remaining genetic variations in current Tibetan populations and no significant differences between haplotypes without the deletion. Besides, this deletion is located close to *PRDM16* (PR/SET domain 16). PRDM16 is a key regulator of the thermogenic program that controls the endogenous transformation of white to brown adipose tissues in response to cold (Seale et al. 2011; Cohen et al. 2014), and has been reported to exhibit up-regulated expression in the Tibetan pig (Lin et al. 2017). In summary, the 335 bp deletion at 1p36.32, which is located in a region with weak positive selection, is likely to be introgressed from the ancient humans and related to the local adaptation of Tibetan highlanders.

**Fig. 6.**
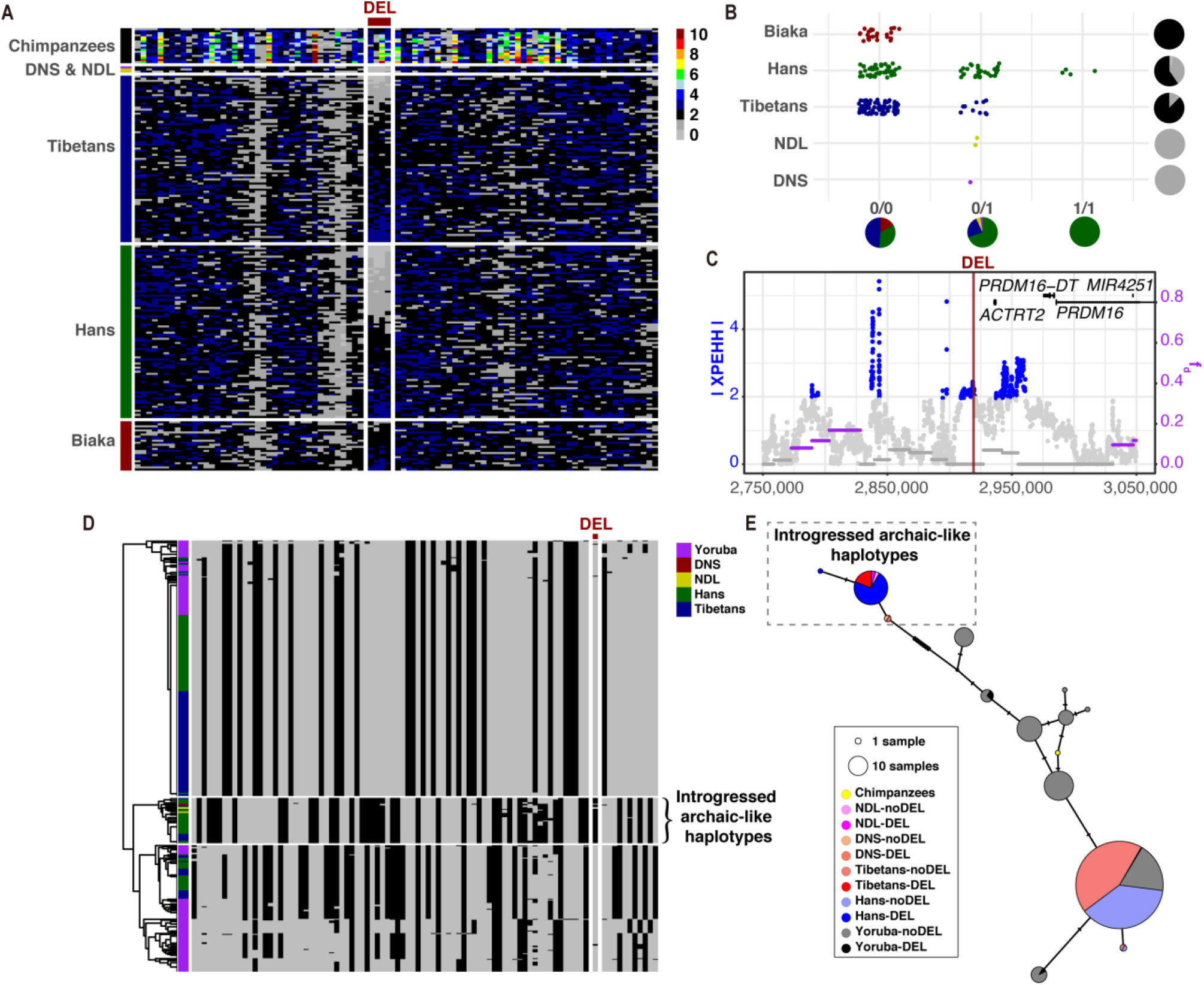
Signatures of selection and archaic introgression from Neandertals (NDL) and Denisovans (DNS) for the deletion of 335 base pairs at 1p36.32. (A) Absolute integer copy numbers for a sliding window of 100 base pairs (bp) and a step size of 50 bp around the deletion. Each row represents the copy numbers of a sample over the region. (B) Genotyping results for the deletion among different populations. Pie charts along the x-axis indicate the population distribution for different structural variation (SV) genotypes (colors are the same as populations), and pie charts along the y-axis illustrate the frequency distribution for a given population (colors are the same as copy number of 2/1/0). (C) Distributions of single nucleotide variants (SNVs) with significant f_d_-statistic (purple dots) and cross-population Extended Haplotype Homozygosity (XP-EHH, blue dots). (D) Hierarchical clustering for the region around the deletion. Rows illustrate the individual haplotypes. The column indicates the genotypes of 90 variants with selection signals (XP-EHH > 2). The color of gray and black represents the ancestral and derived alleles, respectively. (E) TCS haplotype network for 20 linkage disequilibrium (LD)-linked SNVs to the deletion with selection signals (XP-EHH > 2). All haplotypes with the deletion in Tibetans and Hans, and the haplotypes with or without the deletion in the ancient genomes are clustered together within the dotted box.

## Discussion

In this study, we comprehensively analyzed the complete spectrum of SV in a Chinese Tibetan and Han population using the nanopore sequencing technology. We explored novel adaptive genes affected by Tibetan-Han stratified SVs. Besides, we carried out the first study to investigate adaptive introgressed SVs in the whole genome of Tibetans. Overall, our results provide a valuable resource for future HAA studies.

Due to the limitations of the short-read sequencing technology, a substantial amount of SVs remains uninvestigated. Although the publicly available SV calls we used for comparison included both short-read and long-read sequencing data, there are still more than one-quarter of SVs in our catalog that are sequence-resolved for the first time. Therefore, our high-confidence SV callsets provide a considerable number of common variations in Chinese Tibetan and Han populations. Then, we adopted a combined strategy that SVs discovered within a small population using LRS were genotyped with a relatively large amount of NGS data. It is noteworthy that approximately half of the SVs discovered in LRS data can not be recalled in NGS data. Through analyzing the local sequence architecture around SVs, we speculated that the relatively low resolution in the repetitive regions is likely to be the primary defect of SV genotyping with NGS data. Further genotyping studies should focus on improving the unsatisfying recall rates in repetitive regions. To facilitate the use of these resources for the genomics community, we published our callset with high-coverage genotype calls from 189 samples.

Though SVs are known to impact the formation of population diversity and adaptation positively (Moan et al. 2019), few studies have examined SVs’ role in HAA. Considering three types of evidence-supported genes, we screened out a total of 124 protein-coding adaptive genes affected by 90 population-stratified SVs between Tibetans and Hans, providing valuable resources for future HAA studies. Moreover, the majority of these genes have not been reported in previous studies, which could improve our understanding of multi-variant adaptation. For example, *CPEB1* and *ITGA1* are two potential adaptive genes, both of which lead to increased oxygen delivery and promote adaptation in the hypoxic environment (Senger et al. 2002; Hägele et al. 2009). These Tibetan-Han stratified SVs and potential adaptive genes will facilitate further molecular-functional studies of HAA.

Although recent studies have found that genetic variants originated from archaic hominins contribute to local population adaptation (Huerta-Sánchez et al. 2014; Hsieh et al. 2019; Almarri et al. 2020), little is known of adaptive introgressed SVs in Tibetans. We searched for signatures of natural selection and ancient human introgression around Tibetan-Han stratified SVs using simulated neutral variations and finally identified a total of 15 adaptive introgressed candidates. To the best of our knowledge, this is the first study to detect adaptive introgressed SVs in the whole genome of Tibetans. Our results suggested that the introgressed SVs could contribute to the local adaptation of Tibetan highlanders. It should be noted that SVs which were not successfully recalled in any ancient genomes were not necessarily absent from ancient humans, especially considering the short-read of ancient genomes. For instance, there are many flanking SNVs around the 2,590 bp deletion at 7p22.3 that exhibited both signatures of natural selection (|XP-EHH|> 2) and archaic introgression (p < 0.05 for both D-statistic and f_d_-statistic) (Supplemental Fig. S10C; Supplemental Table S13), suggesting that the surrounding genomic region of the deletion could come from archaic introgression and contribute to adaptation. Thus, each candidate needs more detailed investigations in the future.

In conclusion, our study demonstrates that SVs play an important role in the evolutionary processes of Tibetans’ adaptation to the Qinghai-Tibet Plateau. Our comprehensive SV callset, which consists of 38,216 unique SVs, provides a considerable number of previously unresolved common variants in Chinese populations. Furthermore, we screened out a total of 124 protein-coding adaptive genes affected by 90 population-stratified SVs between Tibetans and Hans, which reveals new evidence of HAA and will improve our understanding of multi-variant adaptation. Besides, we carried out the first study to investigate adaptive introgressed SVs in the whole genome of Tibetans and discovered 15 candidates, suggesting that SVs originated from archaic hominins and introgressed into Chinese populations could contribute to the local adaptation of Tibetan highlanders.

## Methods

### Study participants recruitment

We recruited 15 unrelated ethnic Tibetan and 10 unrelated Han subjects for nanopore sequencing (Supplemental Table S1). These 25 subjects were recruited during a physical examination program at the community conducted in December 2018 in Chongqing city located in southern China. Among the 15 indigenous Tibetan subjects, seven came from Shigatse (4,000 meters above sea level), and the others came from areas of relatively lower altitudes (between 3,000 and 4,000 meters). The mean age (s.d.) of these Tibetans is 20.13 (0.99) years old. All Hans came from plain areas (≤ 1,500 meters), and their mean age (s.d.) is 23.20 (4.49) years old. All subjects are males. Besides, we also aggregated several short-read whole-genome sequencing (WGS) datasets (Supplemental Table S2 and Supplemental Methods).

All the above mentioned studies were performed with the approval of the Medical Ethical Committee of the Beijing Institute of Radiation Medicine (Beijing, China). Written informed consent was obtained from each participant.

### Nanopore sequencing

The peripheral blood samples were collected from 25 subjects, and then the genomic DNA was extracted by QIAGEN^®^ Genomic DNA extraction kit (Cat ID: 13323, QIAGEN) according to the standard operating procedure provided by QIAGEN. The extracted DNA was detected by NanoDrop™ One UV-Vis spectrophotometer (Thermo Fisher Scientific, USA) for DNA purity (OD260/280 ranging from 1.8 to 2.0 and OD 260/230 is between 2.0-2.2), then Qubit^®^ 3.0 Fluorometer (Invitrogen, USA) was used to quantify DNA accurately. Since the sample was qualified, BluePippin system (Sage Science, USA) was used to size-select long DNA fragments. Then, we repaired the DNA and prepared the DNA ends for adapter attachment. The sequencing adapters supplied in the SQK-LSK109 kit were attached to the DNA ends. Finally, Qubit^®^ 3.0 Fluorometer (Invitrogen, USA) was used to quantify the size of library fragments. In the end, we primed the Nanopore GridION X5 sequencer (Oxford Nanopore Technologies, UK) flow cell and loaded the DNA library into the flow cell. All samples were sequenced with 1D R9.4.1 nanopores. Each genome was sequenced to a 10~20 × coverage depth.

### SV discovery and genotyping

Base-calling of the raw nanopore squiggles was performed using Guppy v2.3.1 for GridION, and a minimum quality score of 7 (Q7) was applied. Reads were mapped to GRCh37 human reference. Mapping was performed using NGMLR (v0.2.7) (Sedlazeck et al. 2018b) with default parameters for the nanopore sequencing. SV calling was performed using Sniffles (v1.0.11) (Sedlazeck et al. 2018b), NanoSV (v1.2.3) (Stancu et al. 2017), and SVIM (v1.2.0) (Heller and Vingron 2019). Then we systemically assessed the distribution of these SVs across repeat sequences and functional genomic regions. Further information is described in the Supplemental Methods.

SVs discovered using LRS were genotyped with a relatively large amount of NGS data accumulated in previous studies. Reads from all short-read datasets were mapped to GRCh37 human reference. Mapping was performed by BWA-MEM (v0.7.19) (Li 2013) with default parameters. Using the SVs discovered by the nanopore sequencing, we took Paragraph (v2.4a) (Chen et al. 2019) to genotype each genome generated using NGS data (Supplemental Methods).

### Population structure analysis

We calculated the per-individual SV-heterozygosity using VCFtools, which could exhibit relative diversity among populations. PCA and admixture analysis were carried out to infer the covariance structure of allelic frequencies. Next, we employed the Weir and Cockerham estimator for F_ST_ based on VCFtools to identify the Tibetan-Han stratified SVs. We considered three types of evidence-supported genes that tend to be affected by these SVs. Then we used SNVs to estimate the demographic history of Tibetans and Hans, and performed whole-genome coalescent simulations with msprime (Kelleher et al. 2016). Further information is described in the Supplemental Methods.

### Natural selection and archaic introgression

To detect evidence of natural selection in Tibetans, we calculated iHS (Voight et al. 2006) and XP-EHH (Sabeti et al. 2007) using an R package rehh (Gautier and Vitalis 2012). Besides, we applied the D-statistic and f_d_-statistic using admixr (Petr et al. 2019) to distinguish excess genetic drift from the ancient introgression based on allele frequencies. Further information is described in the Supplemental Methods.

## Data access

The basecalled nanopore sequencing data generated in this study have been submitted to the NCBI BioProject database (https://www.ncbi.nlm.nih.gov/bioproject/) under accession number PRJNA681146. All scripts used in this study can be found in the stable release at GitHub (https://github.com/quanc1989/SV-ONT-Tibetan).

## Acknowledgments

This work was supported by the General Program (31771397, 81573251 and 81672369) of the Natural Science Foundation of China (www.nsfc.gov.cn), Beijing Nova Program (20180059), National Key R&D Program of China (No. 2017YFA0504301), and Chinese Key Project for Infectious Diseases (No. 2018ZX10732202 and 2017ZX10203205).

## Author contributions

Conceptualization, C.Q., Y.Lu, and G.Z.; Methodology, C.Q., Y.W., J.P., and Y.Lu; Study participants recruitment, Y.W., J.P., Y.Li, and Y.Lu; Nanopore sequencing data generation and analysis, C.Q., Y.W., and Y.Lu; Illumina NGS data generation and analysis, Y.W, Y.Li, and Y.Lu; Genotyping and population analysis, C.Q., Y.W., and J.P.; Data Curation, C.Q., Y.W., J.P., Y.Li, and Y.Lu; Validation, Y.W. and Y.Li; Manuscript writing, C.Q., Y.Li, Y.Lu, and G.Z; Visualization, C.Q. and Y.Lu; Supervision, Y.Lu and G.Z..

