## Supplemental Material for "Characterization of Structural Variation in Tibetans Reveals New Evidence of High-altitude Adaptation and Introgression"

This file includes:

Supplemental Methods

Supplemental figures S1-S11

Supplemental tables S1, S4-S8, S16 (other tables longer than one page are provided as Excels).

### Supplemental Methods

#### Short-read data aggregation

We used a short-read whole-genome sequencing (WGS) dataset generated by our lab, including 43 unrelated ethnic Tibetans and 46 unrelated individuals of Hans (Supplemental Table S2). These 43 Tibetans were recruited during a physical examination program at the community conducted from June 2010 to August 2010, who live in five counties (including Batang, Dawu, Kangding, Dzongzhab, and Litang) in Sichuan province located in southwest China, at > 2,500 meters over sea level. The male/female ratio and the mean age (s.d.) of these Tibetans are 1.50 and 36.31 (12.31) years old respectively. These 46 Hans were recruited in a regular physical examination from July 2010 to August 2010 at the Guangxi Cancer Hospital (Nanning city, China), who lives in Nanning city, Guangxi province located in southern China, at < 100 meters over sea level. The male/female ratio and the mean age (s.d.) of these Hans are 1.50 and 42.22 (6.39) years old respectively. These samples were sequenced to 150 bp paired-end reads with an average coverage of 40x using the Illumina platform.

In the meantime, we aggregated several short-read WGS datasets published in previous studies. Firstly, we collected a subset of publicly available NGS dataset of 38 Tibetan highlanders and 39 Han lowlanders (Lu et al. 2016). Besides, 24 Biaka in the Central African Republic from the Human Genome Diversity Project (HGDP) was introduced as an outgroup. These samples were also sequenced to 150 bp paired-end reads with an average coverage of 40x. To discover archaic introgression of SVs, we also collected three publicly available archaic hominin genomes, including a Denisovan (Meyer et al. 2012), a Neanderthal (Prüfer et al. 2014) from the Altai Mountains, and another Neanderthal from Croatia (Prüfer et al. 2017). These archaic samples were sequenced to 50 bp paired-end reads with an average coverage of 40x. Chimpanzee WGS data were obtained from Great Ape Genome Project (Prado-Martinez et al. 2013), including 17 individuals which were sequenced to 100 bp paired-end reads with an average coverage of 25x.

#### SV discovery

SV calling was performed using Sniffles (v1.0.11) (Sedlazeck et al. 2018), NanoSV (v1.2.3) (Stancu et al. 2017), and SVIM (v1.2.0) (Heller and Vingron 2019). These tools have been reported to be compatible with NGMLR and show better accuracy and sensitivity than others (Coster et al. 2019). First, we used the tools described above to perform SV detection on each sample. Five minimum supporting reads with at least 50 bp length was required. The insertion sequence and read ID was required for each method, and the rest are all default parameters. SURVIVOR (v1.0.7) (Jeffares et al. 2017) was used to merge the SVs supported by at least two methods with a maximum allowed pairwise distance of 1,000 bp between breakpoints. Meanwhile, the SVs obtained from different tools are not necessary to agree on the SV-type or the strand, so that we could capture as many potential breakpoints as possible. Finally, we merged the SVs obtained from all the samples as long as one sample supports it.

Thus, we have obtained potential regions for all samples, and we need to get a fully genotyped multi-samples dataset. We re-ran Sniffles across all the samples with all these potential regions (--Ivcf) and finally combined the SVs with SURVIVOR. This time, we asked SURVIVOR only to report SVs supported by at least one sample, and they have to agree on the SV-type. Furthermore, we used a hard threshold with five minimum supporting reads, and all non-missing genotypes less than this threshold were modified to reference. To comprehensively understand the local context of SVs, we decided not to filter out centromeric or pericentromeric regions.

**Comparison of SVs to diverse databases**

We compared our SVs to several previously published datasets using AnnotSV (v2.2) (Geoffroy et al. 2018), including the Database of Genomic Variants (DGV), gnomAD (Collins et al. 2020), the Deciphering Developmental Disorders (DDD) Study, the 1000 Genomes Project (Sudmant et al. 2015), and a common disease trait mapping study (Abel et al. 2020). Both short-read and long-read SV calls were included in this dataset. Only the variants overlapping at least 50% of our SVs were reported. We calculated the maximum variant frequencies for SVs found in these databases and annotated each SV with the corresponding database.

**SV distribution**

To measure the SV distribution in the human genome, we divided each chromosome into continuous 500 kb windows and summed the number of SVs overlapped with the window. Windows that had overlapped with centromeres were excluded. For insertions and translocations, we only considered the position of breakpoints. While for inversions, deletions, and duplications, we considered the entire SV region. We calculated the distance from each window’s edge to the nearest telomere, and log2 fold change (LFC) was used to compare the degree of enrichment inside and outside 5 Mb to the telomere. A 100 round permutation test was performed by randomly exchanging the position of windows, and the empirical p-value was calculated by the distribution of LFC for each test.

**Genomic background model**

To perform overlap enrichment analysis of SVs versus genomic elements, we generated 100 randomly shuffled SV sets in the non-gap region of the GRCh37 human genome using BEDTools (v2.28.0) (Quinlan and Hall 2010), as described in the SV release of the 1000 Genomes Project (phase 3) (Sudmant et al. 2015). Each SV in a single shuffled set has the same SV-type and length as the corresponding real SV within the same chromosome but no intersection. These shuffled sets made up a random background model to learn the enrichment of genomic elements overlapped with real SVs versus the null distribution. LFC was used to measure the enrichment, and the empirical p-value was calculated and reported as significant if p-value < 0.05.

**Breakpoint analysis**

AnnotSV was used to identify the overlapping repetitive sequences at the SV breakpoints (+/- 100bp) using RepeatMasker annotations from the University of California Santa Cruz (UCSC) Genome Browser. A breakpoint was annotated as the repeat class of a specific repetitive sequence which intersects with its surrounding sequence. RepeatMasker annotations from the UCSC Genome Browser Database used the RepBase library (Jurka 2000) from the Genetic Information Research Institute (GIRI) and summarized the repetitive sequences into ten different classes, including low complexity repeats, simple repeats, satellite repeats, SINE, LINE, LTR, DNA repeat elements, RNA repeats, other repeats, and unknown ones. We performed permutation tests for each repeat class using the random background model.

Breakpoint junction sequences can be used to infer formation mechanisms for SVs (Lam et al. 2009). We used a simplified pipeline to systematically classify the mechanisms, originally from BreakSeq (Lam et al. 2009) and 1000 Genomes Project (phase 3) (Sudmant et al. 2015). First, SVs were examined for > 50% overlapped by tandem repeats to identify the expansion or contraction of VNTRs. Then, SVs were classified as NAHR if both breakpoints were annotated as the same repeat class. Finally, the SVs overlapped with interspersed repetitive sequences were inferred into TE-mediated mechanisms. We considered only VNTR and TE for insertions.

**Gene annotations and chromatin features**

Consistent with the method described in the GnomAD-SV (Collins et al. 2020), we used the protein-coding gene annotations from Gencode and generated a canonical transcript described in

Ensembl Genomes Project (Hubbard et al. 2008). Then we annotated SVs for a range of potential effects on coding sequences using svtk (Werling et al. 2018), including LoF variants, copy gains (CG), partial gene duplications, whole-gene inversions, intronic, and intergenic SVs. Then we detected the potential purifying selection against gene-altering SVs. Annotations from the Exome Aggregation Consortium (ExAC) (Lek et al. 2016) were used to indicate the probability of a gene intolerance to variations, including synZ (Z score > 0 indicates intolerance to synonymous variations), misZ (Z score > 0 indicates intolerance to missense variations), and pLI (pLI > 0.9 indicates intolerance to loss of function variations). In the meantime, predictions of haploinsufficiency (HI_DDDpercent < 10% indicates more likely to exhibit haploinsufficiency) from DECIPHER (Firth et al. 2009) and assessments of dosage-sensitive (HI_CGscore < 10 indicates more likely to exhibit dosage pathogenicity) from the Clinical Genome Resource (ClinGen) Consortium Rating System were also included. Permutation tests for annotations of all potentially affected genes were performed using the random background model.

Moreover, we used AnnotSV to detect the associations with chromatin boundaries at the borders of TAD and CTCF binding clusters. As described in a previous study (Fudenberg and Pollard 2019), we used BEDTools-cluster on the narrowPeak files from the ENCODE (Dunham et al. 2012) with a merge distance of 5 kb to generate the CTCF binding clusters. To comprehensively characterize the chromatin landscape, we also collected 15-core chromHMM chromatin states applied to 127 epigenomes from RoadMap (Kundaje et al. 2015). Five states were annotated as inactive (8_ZNF/Rpts, 9_Het, 13_ReprPC, 14_PeprPCWk, 15_Quies), and the others were active. Likewise, permutation tests were performed using the random background model.

**SV genotyping and SNV discovery**

As suggested in recent studies (Audano et al. 2019; Coster and Broeckhoven 2019), SVs discovered using LRS were genotyped with a relatively large amount of NGS data accumulated in previous studies. In this way, we could generate a comprehensive SV catalog and meanwhile reduce the average cost. Using the SVs discovered by the nanopore sequencing, we took Paragraph (v2.4a) (Chen et al. 2019) to genotype each genome generated using NGS data. We set the maximum allowed read count for SVs to 20 times the mean genome coverage for each dataset described above. BCFtools (v1.9) was used to merge all genotyped results. Coverage was assessed using mosdepth (v0.2.5) (Pedersen and Quinlan 2017).We also used CNVkit (v0.9.5) (Talevich et al. 2016) to derive the depth-based absolute integer copy numbers for an average bin size of 100 bp around SVs with default thresholds.

In the meantime, we applied the HaplotypeCaller in GATK (v3.7) to call SNVs in each Chinese sample using NGS data. Only the variants that meet the following criteria were retained: (1) SNVs showed only two alleles; (2) the quality score > 20; and (3) SNVs have valid ancestral states derived from the human-chimp alignment published by the 1000 Genomes Project. Then, the genotypes from different panels were combined using BCFtools. SNVs were further phased and imputed by SHAPEIT (v4.1.2) without reference panels (Delaneau et al. 2019).

**Population structure analysis**

We employed the Weir and Cockerham estimator for $F_{ST}$ (Weir and Cockerham 1984) based on VCFtools to identify the Tibetan-Han stratified SVs. We then performed permutation tests by shuffling population labels for 1,000 times and calculated the empirical p-values. We determined a SV is population-stratified in TIB corresponding to HANN or HANS if (1) $F_{ST} > 0.1$, (2) empirical p-value < 0.05, and (3) missing rate < 0.1. In order to identify all candidate adaptive genes that tend to be regulated by these SVs, we considered three types of evidence-supported genes, including linear overlapping/nearest genes, 3D affected genes, and LD-linked eGenes.

**LD-linked eQTLs**

To study the potential effects of SVs on gene expression, we extracted all significant variant-gene pairs of 49 tissues from the Genotype-Tissue Expression (GTEx) project (v8 release) (GTEx Consortium 2020). Meanwhile, we used causal posterior probability (CPP) of fine-mapping results from CAVIAR to identify the causal variants(Hormozdiari et al. 2014; Chiang et al. 2017). eQTLs with the most significant variants for genes or CPP > 0.1 were reserved for the following analysis. Then, we tested whether these casual variants and SVs were in LD ($\pm$1 Mb, $r^2 > 0.8$). LD association tests were performed with VCFtools using both Tibetan and Han samples. Next, we identified the potentially regulated genes from LD-linked SV eQTLs and created a body map by Gene ORGANizer (Gokhman et al. 2017). We also plotted the heatmaps of associated eQTLs across genes by LDheatmap (Shin et al. 2006).

**Chromatin conformational alterations**

We applied a recently published algorithm to model the chromatin conformational changes caused by population-stratified SVs (Sadowski et al. 2019; Wlasnowolski et al. 2020). Briefly, deletions can remove all overlapped anchors, and insertions will construct new CTCF anchors connected with the closest anchors if CTCF motifs were identified in the inserted sequences. We collected clusters of CTCF and RNA polymerase II derived from Chromatin Interaction Analysis by Paired-End Tag Sequencing (ChIA-PET) experiments and Hi-C maps of GM12878 from previous studies (Rao et al. 2015; Sadowski et al. 2019). Circular layouts were generated for the visualization of altered interactions by circlize package in R (Gu et al. 2014).

**Enrichment analysis**

We predicted the effect of population-stratified SVs on transcripts and regulatory regions in the linear genome by the Ensembl Variant Effect Predictor (McLaren et al. 2016). Combined with the LD-linked genes and predicted affected genes in the 3D genome, we discovered a total of 124 protein-coding genes. Enriched biologically functional annotations of these candidate adaptive genes were analyzed by Metascape (Zhou et al. 2019). Besides, we collected the hypoxia-related pathways from a recent study (Deng et al. 2019) and expanded the list of known adaptive genes through literature analysis (Supplemental Table S14). Fisher's exact test was performed to test the pathway enrichment.

**Demographic inference**

As suggested in a recent study (Hsieh et al. 2019), we used SNVs to estimate the demographic history of Tibetans and Hans. SNVs were excluded if they overlapped with any low complexity regions in RepeatMasker annotations or self-chains (sequence identity > 0.9) in the UCSC Genome Browser Database. As suggested in previous reports (Hsieh et al. 2016; Hsieh et al. 2019; Liu et al. 2019), the coding sequences (Refseq genes from the UCSC Genome Browser Database) with 1,000 flanking base pairs were excluded to avoid potential effects of natural selection. All SNVs were pruned using PLINK (v1.9) in a window of 1,000 variants with a step size of 5 and a pairwise $r^2$ threshold of 0.5. At last, an unfolded joint site frequency spectrum (SFS) of 876,639 SNVs from 160,656,471 non-genic autosomal bases was estimated using easySFS (https://github.com/isaacovercast/easySFS), polarized using ancestral alleles.

In order to explore the alternative demographic models for the population model TIB-HANN-HANS, we used the diffusion approximation method of ∂a∂i (Gutenkunst et al. 2009) to analyze the joint SFS. We tested three models with symmetric migrations to describe the simplified evolution paths. All three populations differentiated at the same time in the first model, while TIB branched out first in the other two models. Additional parameters were added to the third model to describe the possible changes in population size. To each model we tested, consecutive nine rounds of optimizations (100 replicates) were performed following the dadi_pipeline workflow (Portik et al. 2017). For each round, we ran multiple replicates and used parameter estimates from the highest log-likelihood replicate to seed searches in the following round and optimized parameters using the derivative-based BFGS algorithm. To estimate the parameter uncertainties, we generated 100 nonparametric bootstrap replicates from the whole-genome data and estimated the confidence intervals for parameters using the Godambe information matrix (Coffman et al. 2015). All the scaled parameter estimates were transformed into real values using a mutation rate of $1.5 \times 10^{-8}$ per site per generation (Scally 2016) and a generation time of 29 years (Hsieh et al. 2019).

To discover the potential adaptive introgression from hominins into Tibetans, we grouped SNV genotypes of HANN and HANS into one cluster and then performed demographic inference for another population model YRI-TIB-HAN. We downloaded the SNV genotypes for YRI from the 1000 Genomes Project (phase 3). Using the same pipeline described above, we analyzed another unfolded joint SFS of 1,702,937 SNVs from 174,495,893 non-genic autosomal bases.

**Coalescent simulations**

When the best-fit demographic model was recognized, we used msprime (Kelleher et al. 2016) to perform whole-genome coalescent simulations. To approximately account for mutational heterogeneity across the genome, we applied a three-step framework described in a previous study (Hsieh et al. 2016). First, we divided the genome into windows of 50 kb and used ∂a∂i to estimate the population genetic mutation parameter θ given the best-fit demographic model. Second, we performed a whole-genome simulation with the $\theta_{max}$ estimated among all the windows. The HapMap recombination map (Frazer et al. 2007) was also incorporated to model the recombination heterogeneity. Third, we changed the local mutation rate for each window by randomly dropping a proportion of $1 - \frac{\theta_j}{\theta_{max}}$ of the simulated neutral variants. To account for the parameter uncertainties, we simulated 1,000 replicates of the best-fit demographic model by randomly sampling values from the confidence intervals of each parameter, assuming that they had a multivariate normal distribution. When simulating the best-fit demographic model for YRI-TIB-HAN, we included the Chimpanzee (n = 1) and archaic hominins (n = 3) branches as suggested in a recent study (Hsieh et al. 2019). Genotypes for archaic genomes were collected via Neandertal and Denisovan Genome Projects. Confidence intervals for these relevant parameters were also drawn from this study.

At last, we estimated the per-base mutation rate using Watterson Estimator (Watterson 1975). The significant correlation between the simulated and real data suggested that our whole-genome coalescent simulations could recapitulate the mutation patterns of SNVs. Furthermore, the genomic distribution of $F_{ST}$ was remarkably different between the simulated and real data, indicating that the simulated data could identify non-neutral variants.

**Natural selection and archaic introgression**

To detect evidence of natural selection in Tibetans, we calculated iHS (Voight et al. 2006) and XP-EHH (Sabeti et al. 2007) using an R package rehh (Gautier and Vitalis 2012). We detected selection signals for a sliding window of 100 SNVs and a step size of 50 SNVs, and then determined selection candidates if they were flanked by a significant window (p-value < 0.05, ±500 kb) exclusively for Tibetans. The p-value of each window was defined as the proportion of simulations (TIB-HANN-HANS) with statistic values greater than those in the real data.

To detect evidence of archaic introgression, we applied the D-statistic and $f_d$-statistic using admixr (Petr et al. 2019). D-statistic and $f_d$-statistic are designed to distinguish excess genetic drift from the ancient introgression based on allele frequencies of four populations. Following the instructions of a previous study (Hsieh et al. 2019), we defined the population relationships among three populations and an outgroup to be (((P1, P2), P3), O) = (((YRI, TIB), ARC), Chimpanzee), where ARC represents the Denisovan (DNS, n = 1) or Neanderthal (NDL, n = 2). Different from the relatively short window used for detecting natural selection, we applied the D-statistic and $f_d$-statistic for a sliding window of 500 SNVs and a step size of 250 SNVs. Besides, at least 100 valid SNVs were required for each window. Then, we determined the candidate introgressed SVs if they were flanked by a significant window (p-value < 0.05 for both D-statistic and $f_d$-statistic, ±500 kb) for Tibetans, regardless of Hans. The simulations of the population model YRI-TIB-HAN was regarded as the null distributions in this step. In the meantime, we performed hierarchical clustering for haplotypes by the R function 'hclust' with pairwise nucleotide differences as the input matrix. The TCS haplotype network was constructed using PopART (Leigh and Bryant 2015).

### Supplemental Figures

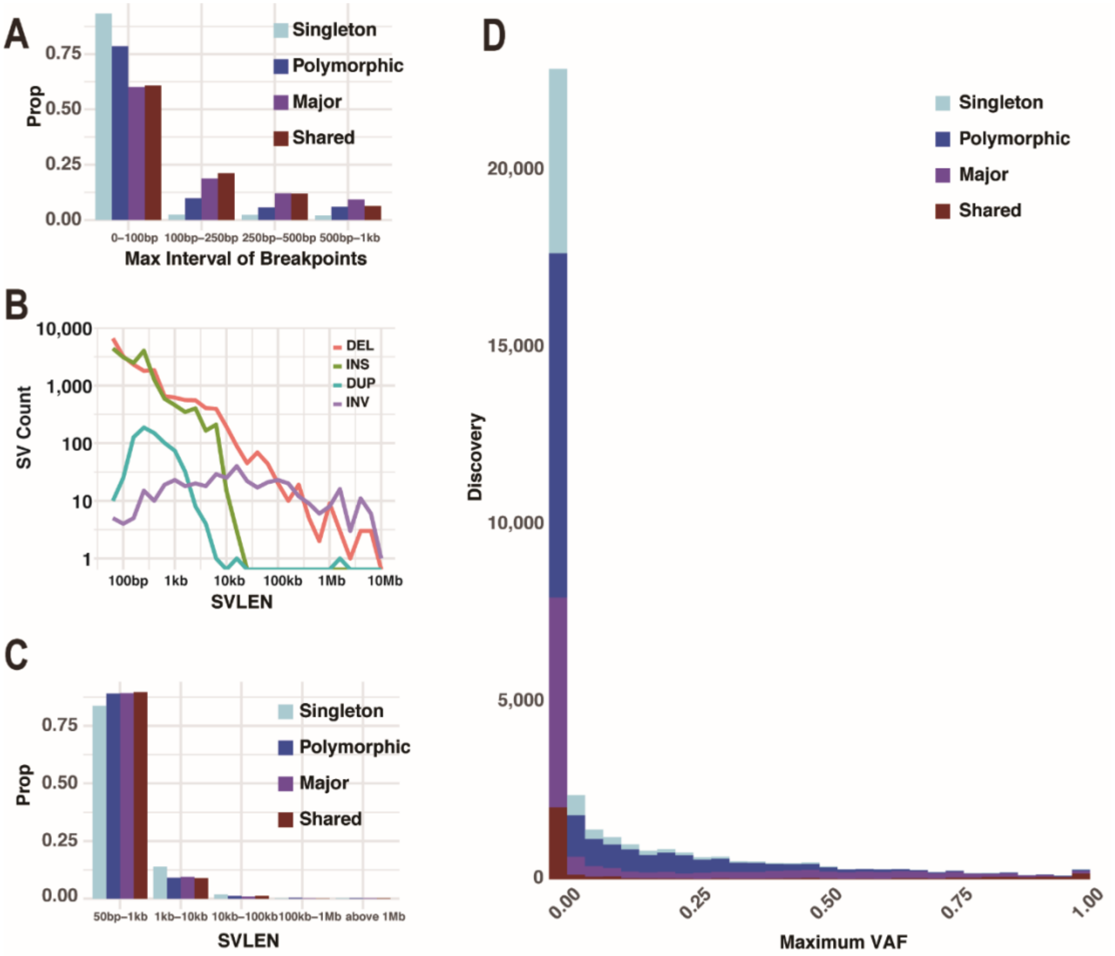


**Fig. S1 Discovery of structural variations in 25 samples using the nanopore sequencing technology. (**A) Maximum interval of breakpoints determined in different samples for each discovery category, including the shared (identified in all samples), major (identified in ≥ 50% of samples), polymorphic (identified in > 1 sample), and singleton (identified in only one sample) structural variations (SVs). (B) Distribution of SV length (SVLEN) for each SV type: inversion (INV), duplication (DUP), insertion (INS), and deletion (DEL). (C) Distribution of SV length (SVLEN) for each discovery category. (D) The maximum variant allele frequencies (VAF) reported in previously published SV calls for each discovery category.

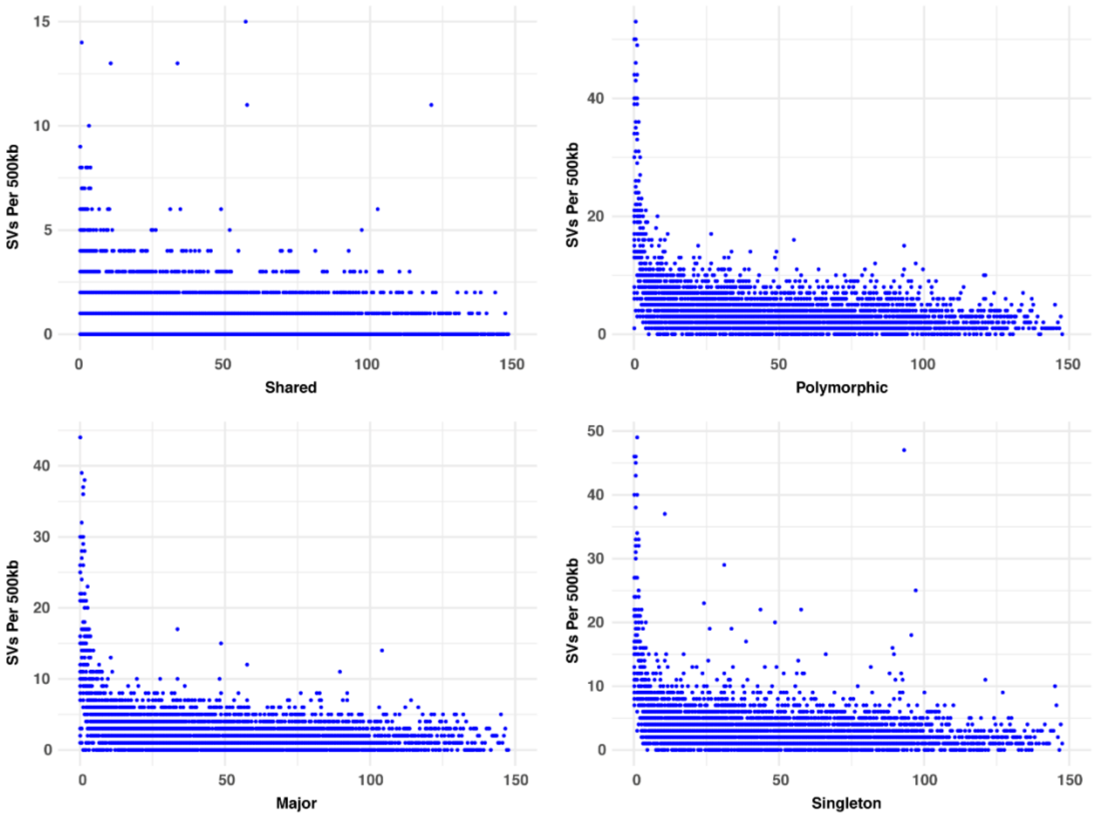


**Fig. S2 Distribution of structural variations.** Each chromosome was divided into continuous 500 kilobase (kb) windows and the number of structural variations (SVs) overlapped with the window was summed for each discovery category, including the shared (identified in all samples), major (identified in ≥ 50% of samples), polymorphic (identified in > 1 sample), and singleton (identified in only one sample) SVs.

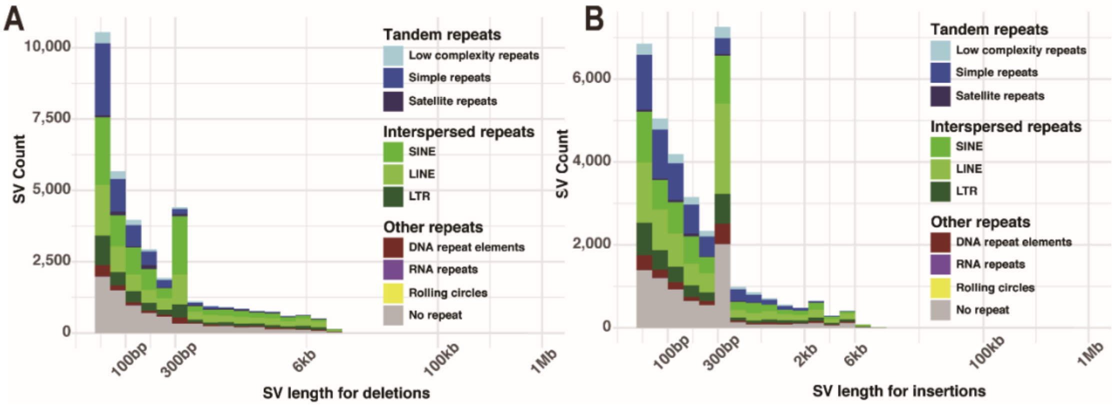


**Fig. S3 Distribution of deletions and insertions classified by intersected repeat elements.** Repetitive elements are summarized into nine different classes, including the low complexity repeats, simple repeats, satellite repeats, short (SINE) and long interspersed nuclear elements (LINE), long terminal repeat elements (LTR), DNA repeat elements, RNA repeats, and Rolling circles.

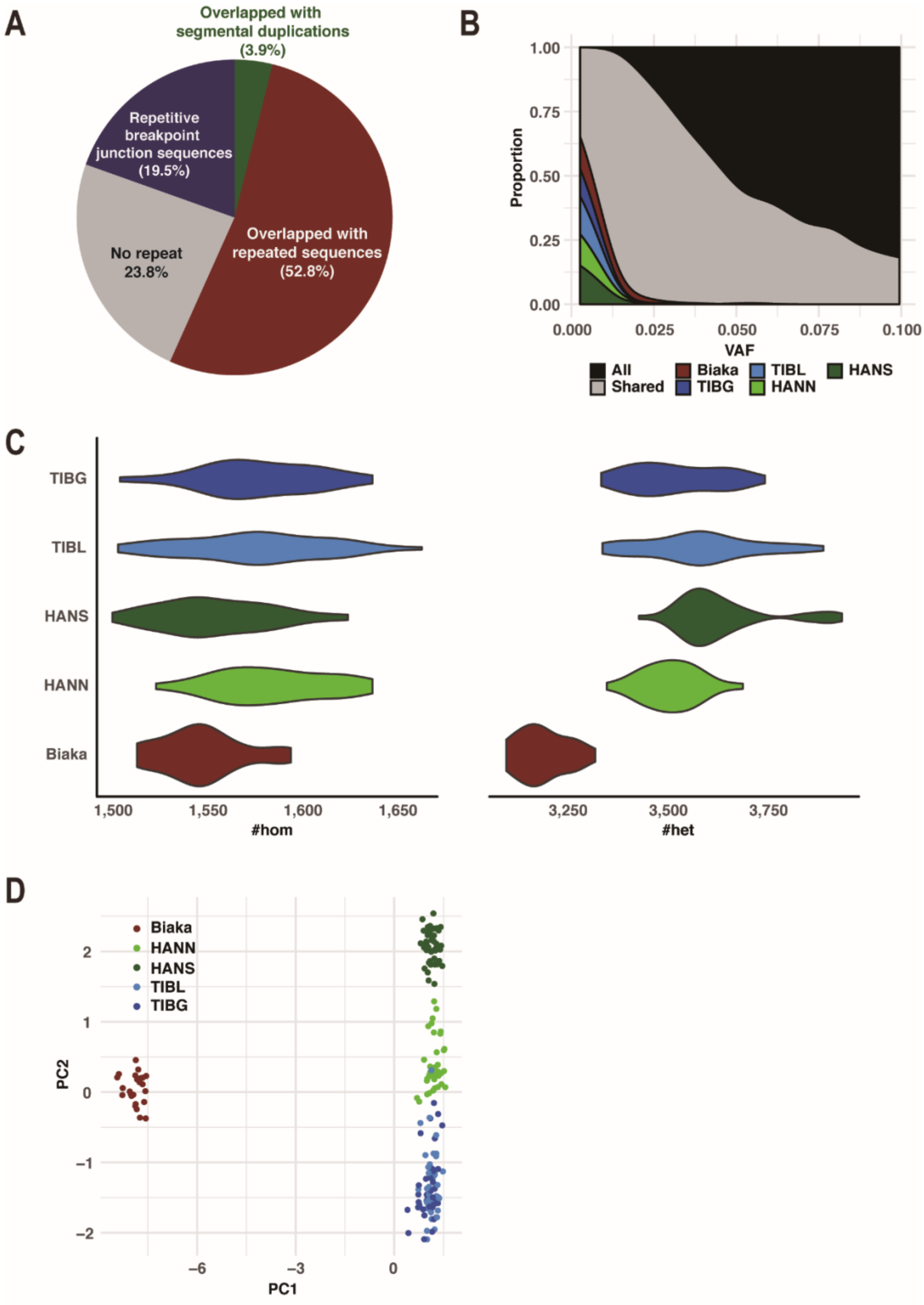


**Fig. S4 Genotyping results and population structure of short-read sequencing genomes.**

(A) Proportions for repeat elements intersected with structural variations (SVs) which were not supported by any next-generation sequencing (NGS) genomes. (B) Profiles of variant allele frequencies (VAF) for each population, including Hans in North China (HANN) or South China (HANS), Tibetans living above (TIBG) or below (TIBL) 4,000 meters, and Biaka populations. (C) Per-individual SV-heterozygosity (#het) and homozygosity (#hom) for each population. (D) Principle component analysis (PCA) of SV genotypes for Biaka, Tibetans, and Hans.

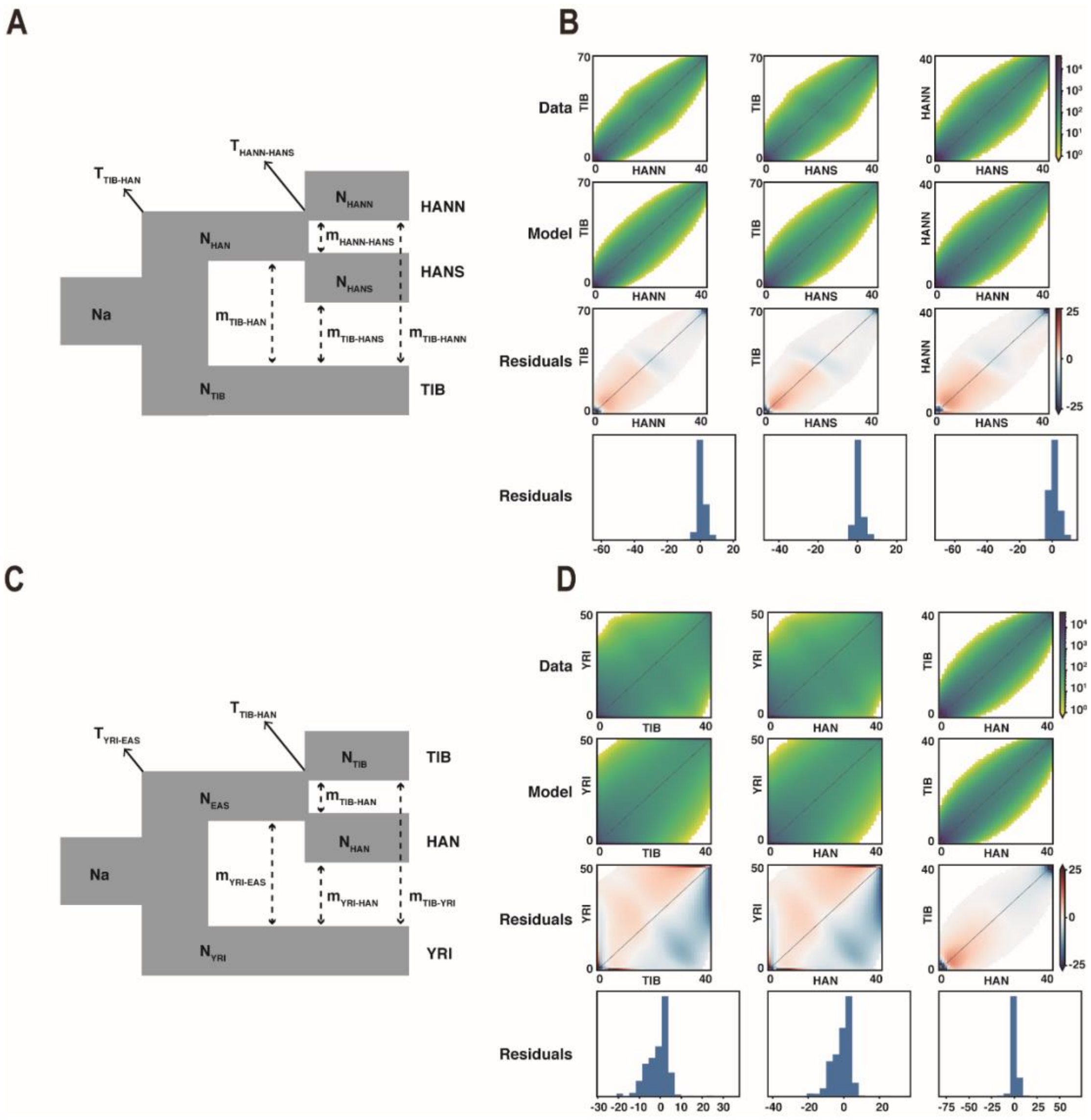


**Fig. S5 Demographic inferences for the best-fit demographic models. (**A) The best-fit demographic model for Tibetans (TIB), Hans in North China (HANN) and South China (HANS). (C) The best-fit demographic model for Tibetans (TIB), Hans (HAN), and Yoruba samples in Ibadan (YRI). The corresponding maximum likelihood estimates and descriptions of parameters can be found in Supplemental Table 7. (B) and (D) The observed and predicted frequency spectra for the two population models, respectively. The first row is the real data, and the second row is the best-fit model. The third and the fourth row are the residuals of the model minus data.

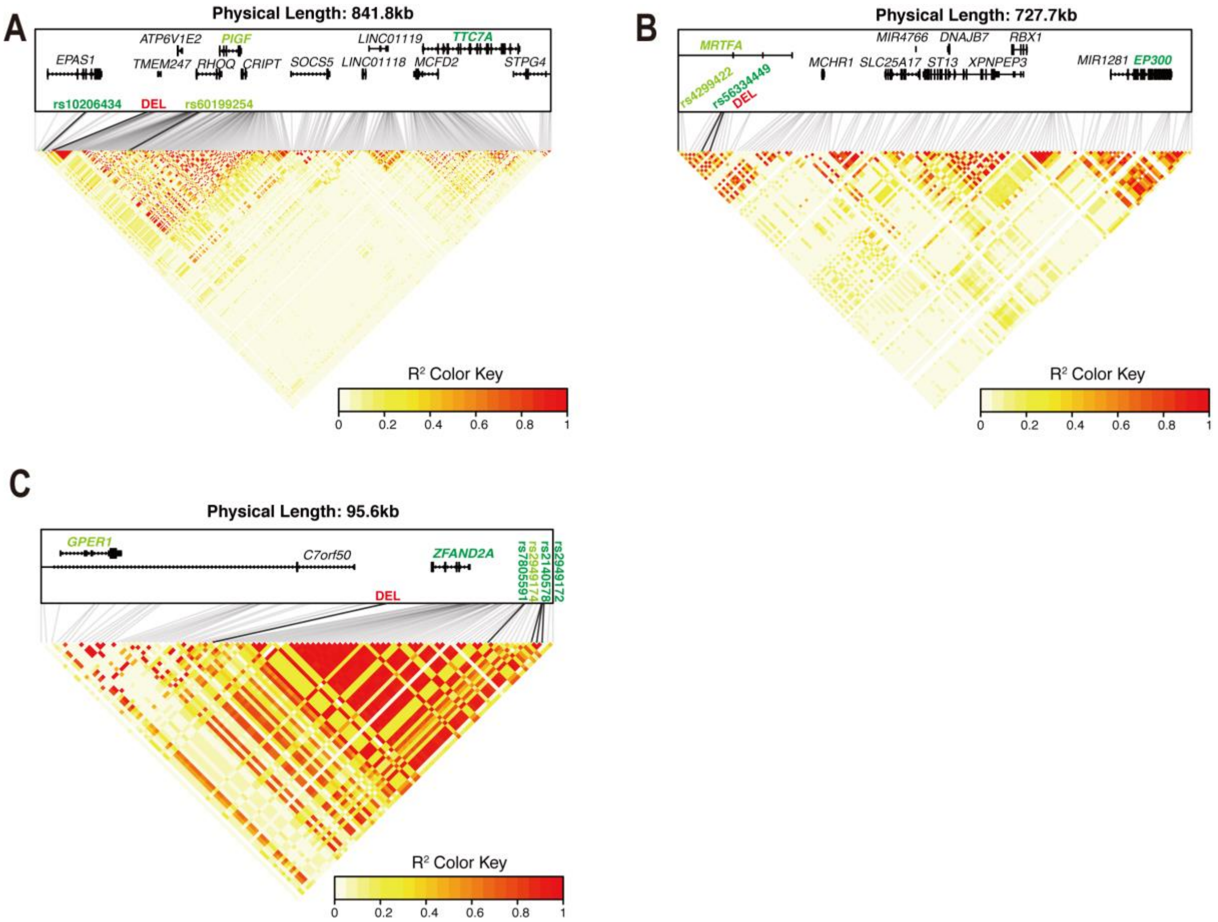


**Fig. S6 Heatmaps for the expression quantitative trait locus (eQTLs) surrounding three deletions, including** (A) the 3.4 kilobase (kb) Tibetan enriched deletion (TED), (B) the 163 bp deletion near *MRTFA*, and (C) the 2,590 bp deletion between *ZFAND2A* and *GPER1*. The linkage disequilibrium (LD)-linked eQTLs and their associated genes are marked as green.

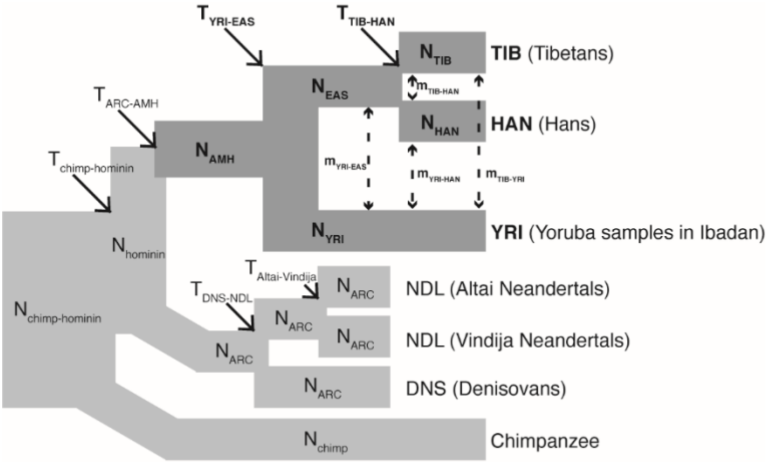


**Fig. S7 Demographic model for the whole-genome coalescent simulations for Tibetans, Hans and Yoruba samples in Ibadan.** Dark branches and bold parameters indicate the best-fit demographic model inferred in this study (Table S7). Parameter estimates of the light branches were uniformly drawn from the 95% confidence interval (CI) reported in a previous study (Table S8).

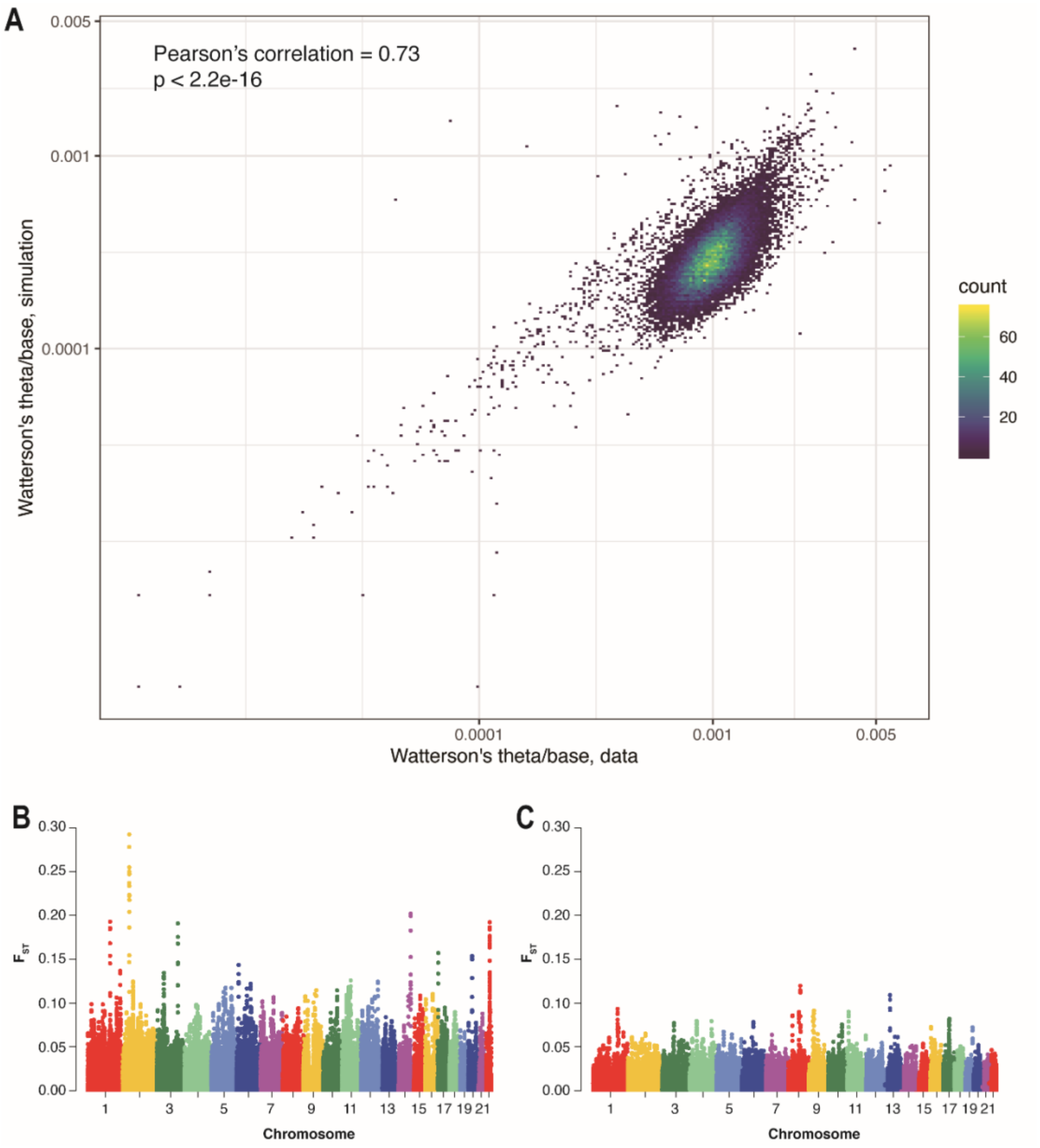


**Fig. S8 Whole-genome simulations for Tibetans, Hans in North China and South China.** (A) Correlation of per-base Watterson Estimator between the simulated and real data. (B) and (C) The Manhattan plots for the window-based $F_{ST}$ statistics (Tibetans *vs.* Hans) using the real and simulated data based on one of 1,000 replicates.

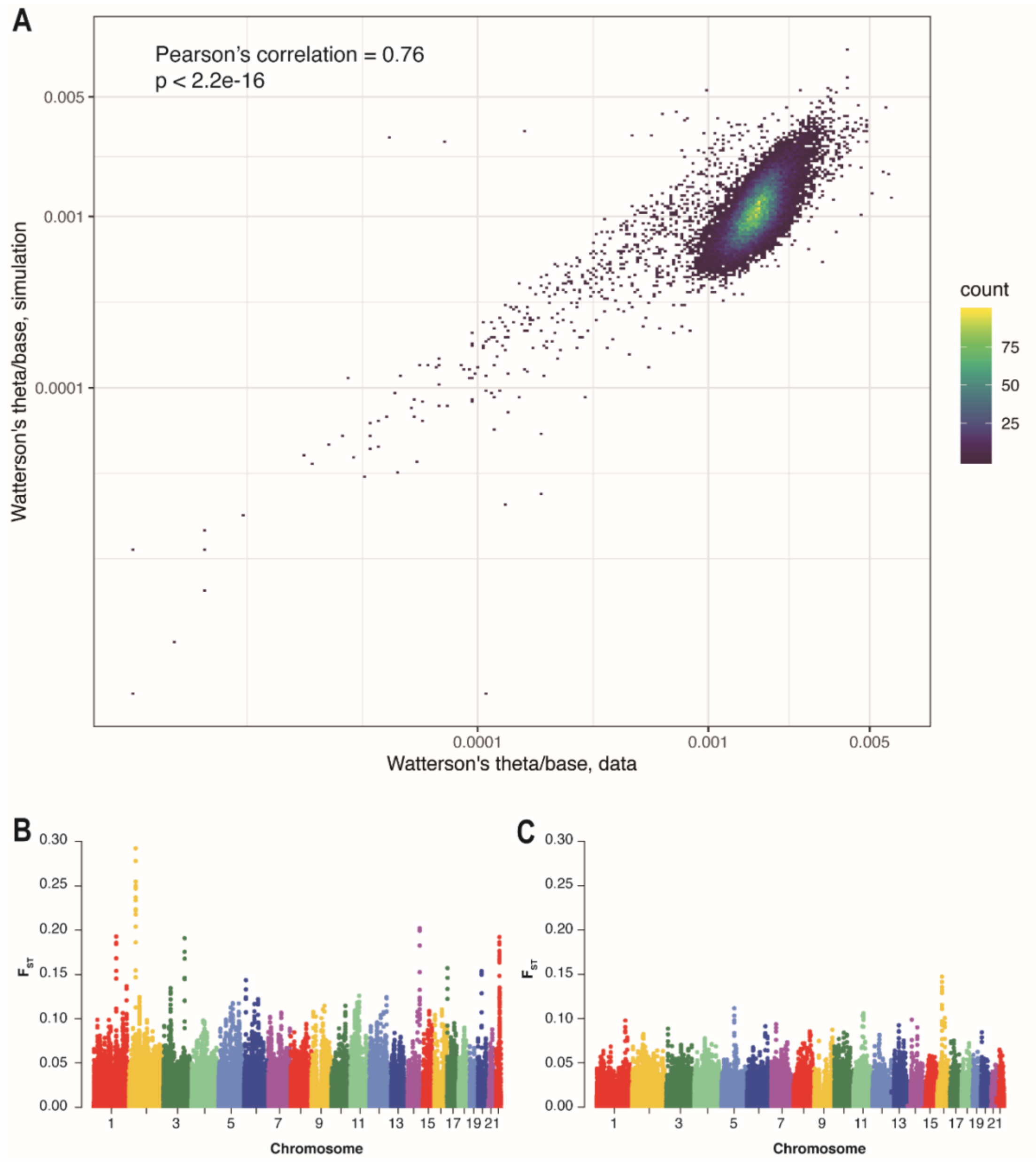


**Fig. S9 Whole-genome simulations for Tibetans, Hans and Yoruba samples in Ibadan.** (A) Correlation of per-base Watterson Estimator between the simulated and real data. (B) and (C) The Manhattan plots for the window-based $F_{ST}$ statistics (Tibetans *vs.* Hans) using the real and simulated data based on one of 1,000 replicates.

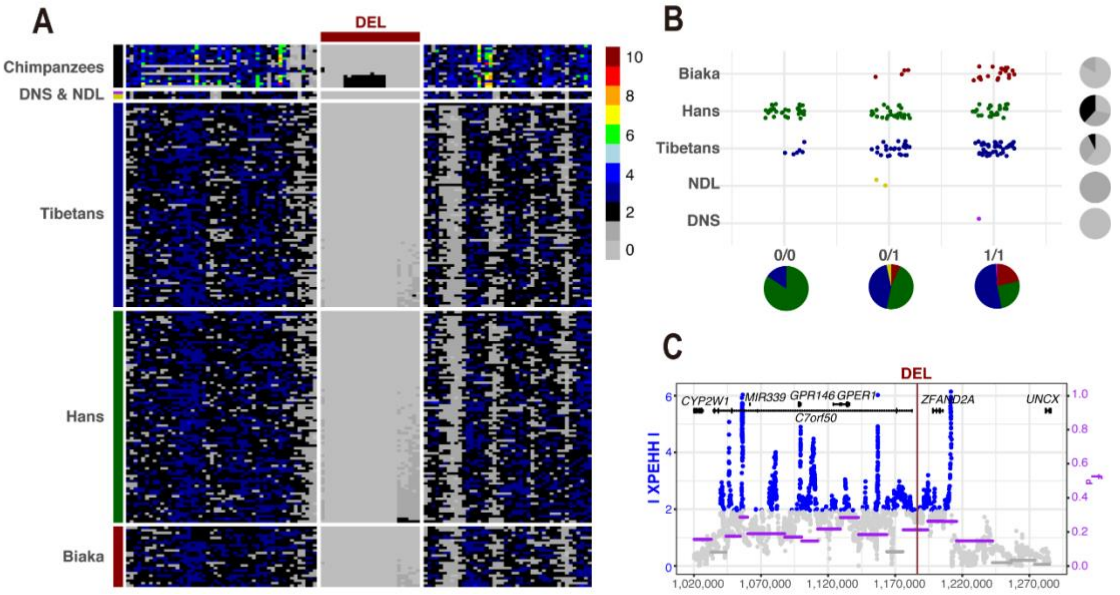


**Fig. S10 Signatures of selection and archaic introgression from Neandertals (NDL) and Denisovans (DNS) for the deletion at 7p22.3 (chr7:1,185,070-1,187,063).** (A) Absolute integer copy numbers for a sliding window of 100 base pairs (bp) and a step size of 50 bp around the deletion. Each row represents the copy numbers of a sample over the region. (B) Genotyping results. Pie charts along the x-axis indicate the population distribution for different structural variation (SV) genotypes (colors are the same as populations), and pie charts along the y-axis illustrate the frequency distribution for a given population (colors are the same as copy number of 2/1/0). (C) Distributions of single nucleotide polymorphisms (SNPs) with significant $f_d$-statistic (purple dots) and cross-population Extended Haplotype Homozygosity (XP-EHH, blue dots).

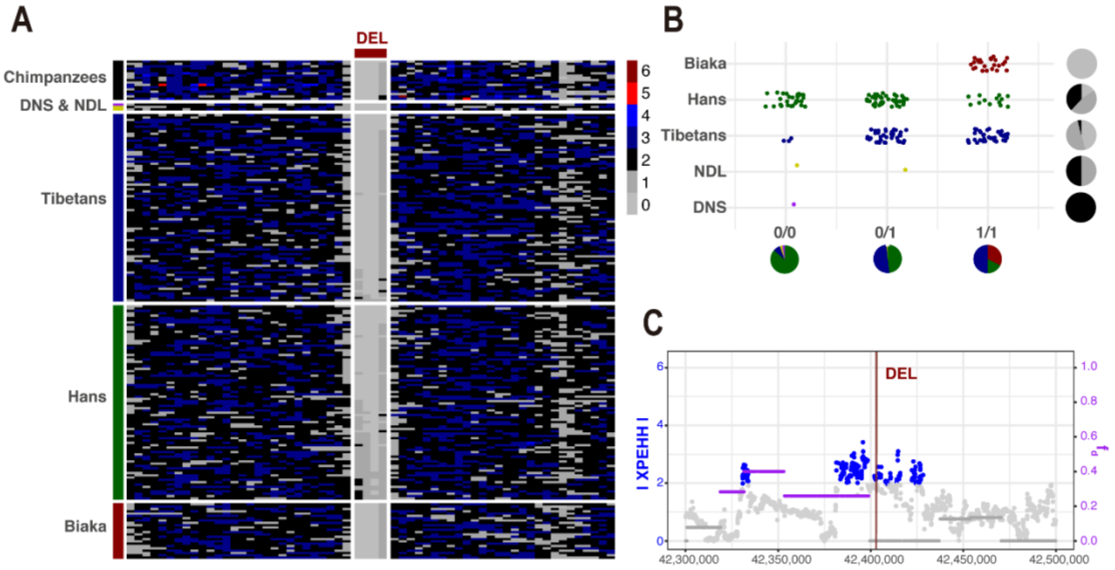


**Fig. S11 Signatures of selection and archaic introgression from Neandertals (NDL) and Denisovans (DNS) for the deletion at 21q22.2 (chr21:42,402,890-42,403,212).** (A) Absolute integer copy numbers for a sliding window of 100 base pairs (bp) and a step size of 50 bp around the deletion. Each row represents the copy numbers of a sample over the region. (B) Genotyping results. Pie charts along the x-axis indicate the population distribution for different structural variation (SV) genotypes (colors are the same as populations), and pie charts along the y-axis illustrate the frequency distribution for a given population (colors are the same as copy number of 2/1/0). (C) Distributions of single nucleotide polymorphisms (SNPs) with significant $f_d$-statistic (purple dots) and cross-population Extended Haplotype Homozygosity (XP-EHH, blue dots).

### Supplemental Tables

**Table S1. Sample information for SV discovery.**

| SampleID | Platform | Populations | Locations | Altitude (meters) | Depth of coverage |
|---|---|---|---|---|---|
| CQ082 | ONT | Tibetan | Hainanzhou | 3,000 | 21.83 |
| CQ085 | ONT | Tibetan | Shigatse | 4,000 | 16.76 |
| CQ091 | ONT | Tibetan | Chamdo | 3,200 | 26.97 |
| CQ095 | ONT | Tibetan | Shigatse | 4,000 | 18.13 |
| CQ104 | ONT | Tibetan | Shigatse | 4,000 | 19.22 |
| CQ115 | ONT | Tibetan | Shigatse | 4,000 | 19.65 |
| CQ121 | ONT | Tibetan | Nyingchi | 3,000 | 13.91 |
| CQ123 | ONT | Tibetan | Hainanzhou | 3,000 | 19.35 |
| CQ132 | ONT | Tibetan | Chamdo | 3,200 | 19.47 |
| CQ136 | ONT | Tibetan | Shigatse | 4,000 | 16.39 |
| CQ143 | ONT | Tibetan | Hainanzhou | 3,000 | 22.97 |
| CQ144 | ONT | Tibetan | Hainanzhou | 3,000 | 20.29 |
| CQ152 | ONT | Tibetan | Hainanzhou | 3,000 | 24.75 |
| CQ188 | ONT | Tibetan | Shigatse | 4,000 | 21.55 |
| CQ200 | ONT | Tibetan | Shigatse | 4,000 | 14.45 |
| CQ232 | ONT | Han | Bijie | 1,500 | 29.30 |
| CQ233 | ONT | Han | Kunming | 1,500 | 11.61 |
| CQ234 | ONT | Han | Zigong | 300 | 27.13 |
| CQ235 | ONT | Han | Lishui | 300 | 24.64 |
| CQ236 | ONT | Han | Ganzhou | 100 | 21.93 |
| CQ292 | ONT | Han | Fuzhou (Jiangxi) | 100 | 15.43 |
| CQ294 | ONT | Han | Panzhihua | 1,000 | 16.03 |
| CQ295 | ONT | Han | Wuyuan | 100 | 13.24 |
| CQ296 | ONT | Han | Ya'an | 500 | 18.51 |
| CQ297 | ONT | Han | Zhengzhou | 100 | 25.96 |

Abbreviations: ONT, Oxford nanopore technologies; SV, structural variation

**Table S4. Enrichment analysis results.**

| Characteristics | Description | LFC | p-value | sd |
|---|---|---|---|---|
| SD | Segmental duplications | 0.168 | 0.010 | 0.057 |
| CpG | CpG islands | 1.866 | 0.010 | 0.239 |
| TAD | Topologically associating domains | -0.040 | 0.000 | 0.018 |
| CTCF | CTCF binding clusters | -0.546 | 0.000 | 0.106 |
| RepClass: Low complexity repeats | Tandem repeats | 1.160 | 0.010 | 0.124 |
| RepClass: Simple repeats | Tandem repeats | 3.419 | 0.010 | 0.357 |
| RepClass: Satellite repeats | Tandem repeats | 2.177 | 0.010 | 0.246 |
| RepClass: SINE | Short interspersed nuclear element | 0.294 | 0.010 | 0.033 |
| RepClass: LINE | Long interspersed nuclear element | -0.357 | 0.000 | 0.037 |
| RepClass: LTR | Long terminal repeats | -0.318 | 0.000 | 0.035 |
| RepClass: DNA | DNA repeat elements | -0.552 | 0.000 | 0.060 |
| RepClass: RNA | RNA repeat elements | -0.155 | 0.168 | 0.166 |
| RepClass: RC | Rolling circles | -1.395 | 0.000 | 0.366 |
| RepClass: No repeats | | -0.607 | 0.000 | 0.062 |
| SVTK: UTR | Untranslated regions | -1.766 | 0.000 | 0.205 |
| SVTK: DUP_PARTIAL | Patial-gene duplications | -0.019 | 0.386 | 0.142 |
| SVTK: LOF | Loss-of-function variants | -1.887 | 0.000 | 0.201 |
| SVTK: INV_SPAN | Whole-gene inversions | 0.551 | 0.119 | 0.537 |
| SVTK: INTRONIC | Intronic SVs | -0.012 | 0.208 | 0.013 |
| SVTK: INTERGENIC | Intergenic SVs | 0.079 | 0.010 | 0.013 |
| ExAC: synZ | Intolerance to synonymous variations | -0.015 | 0.089 | 0.013 |
| ExAC: misZ | Intolerance to missense variations | -0.016 | 0.050 | 0.010 |
| ExAC: pLI | Intolerance to loss of function variations | -0.174 | 0.000 | 0.028 |
| Dosage-sensitive: HI_CGscore | haploinsufficiency | -0.114 | 0.010 | 0.061 |
| Dosage-sensitive: HI_DDDpercent | dosage pathogenicity | -0.433 | 0.000 | 0.049 |
| ChromHMM: 1_TssA | Active TSS | -0.279 | 0.000 | 0.109 |
| ChromHMM: 2_TssAFlnk | Flanking Active TSS | -1.356 | 0.000 | 0.155 |
| ChromHMM: 3_TxFlnk | Transcr. at gene 5' and 3' | -1.288 | 0.000 | 0.238 |
| ChromHMM: 4_Tx | Strong transcription | -0.552 | 0.000 | 0.070 |
| ChromHMM: 5_TxWk | Weak transcription | 0.146 | 0.010 | 0.022 |
| ChromHMM: 6_EnhG | Genic enhancers | -0.928 | 0.000 | 0.112 |
| ChromHMM: 7_Enh | Enhancers | -0.891 | 0.000 | 0.094 |
| ChromHMM: 8_ZNF | ZNF genes & repeats | 0.883 | 0.010 | 0.158 |
| ChromHMM: 9_Het | Heterochromatin | 0.564 | 0.010 | 0.069 |
| ChromHMM: 10_TssBiv | Bivalent/Poised TSS | 0.031 | 0.198 | 0.146 |
| ChromHMM: 11_BivFlnk | Flanking Bivalent TSS/Enh | -0.279 | 0.099 | 0.164 |
| ChromHMM: 12_EnhBiv | Bivalent Enhancer | -0.501 | 0.000 | 0.116 |
| ChromHMM: 13_ReprPC | Repressed PolyComb | 0.018 | 0.386 | 0.067 |
| ChromHMM: 14_ReprPCWk | Weak Repressed PolyComb | 0.403 | 0.010 | 0.045 |
| ChromHMM: 15_Quies | Quiescent/Low | -0.055 | 0.000 | 0.008 |

**Table S5. Comparison of three demographic models for TIB-HANN-HANS.**

| Model ID | Model | Description | Parameter: estimate | | | LL | AIC |
|---|---|---|---|---|---|---|---|
| | | | Effective population size | Divergence time (years) | Migration rate | | |
| Model 1<br>(7 parameters) | 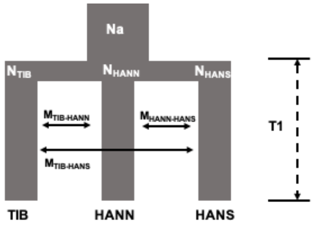 | All three populations differentiated at the same time, including Tibetans (TIB), Hans in the North (HANN), and Hans in the South (HANS). | Na: 14,582<br>$N_{TIB}$: 33,815<br>$N_{HANN}$: 72,561<br>$N_{HANS}$: 31,949 | T1: 14,382 | $M_{TIB\text{-}HANN}$: 0.0606<br>$M_{HANN\text{-}HANS}$: 0.0013<br>$M_{TIB\text{-}HANS}$: 0.0089 | -162,142 | 324,298 |
| Model 2<br>(10 parameters) | 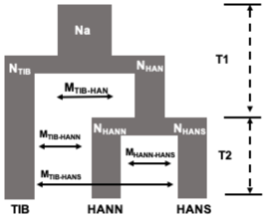 | TIB branched out first | Na: 10,206<br>$N_{TIB}$: 36,260<br>$N_{HAN}$: 50,870<br>$N_{HANN}$: 50,667<br>$N_{HANS}$: 1,632 | T1: 16,871<br>T2: 769 | $M_{TIB\text{-}HAN}$: 0.0016<br>$M_{TIB\text{-}HANN}$: 0.0087<br>$M_{HANN\text{-}HANS}$: 0.0537<br>$M_{TIB\text{-}HANS}$: 0.003 | -152,792 | 305,605 |
| Model 3<br>(17 parameters) | 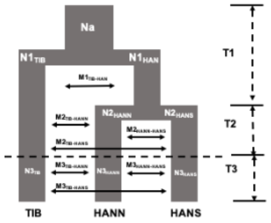 | TIB branched out first and population size changed | Na: 10,853<br>$N1_{TIB}$: 43,328<br>$N1_{HAN}$: 14,389<br>$N2_{HANN}$: 50,667<br>$N2_{HANS}$: 6,318<br>$N3_{TIB}$: 21,323<br>$N3_{HANN}$: 54,195<br>$N3_{HANS}$: 32,581 | T1: 3,147<br>T2: 1,133<br>T3: 9,442 | $M1_{TIB\text{-}HAN}$: 0.0018<br>$M2_{TIB\text{-}HANN}$: 0.0085<br>$M2_{HANN\text{-}HANS}$: 0.081<br>$M2_{TIB\text{-}HANS}$: 0.0058<br>$M3_{TIB\text{-}HANN}$: 0.0988<br>$M3_{HANN\text{-}HANS}$: 0.001<br>$M3_{TIB\text{-}HANS}$: 0.0722 | -160,563 | 321,161 |

LL, log likelihood; AIC, Akaike information criterion.

**Table S6. Comparison of three demographic models for YRI-TIB-HAN.**

| Model ID | Model | Description | Parameter: estimate | | | LL | AIC |
|---|---|---|---|---|---|---|---|
| | | | Effective population size | Divergence time (years) | Migration rate | | |
| Model 1<br>(7 parameters) | 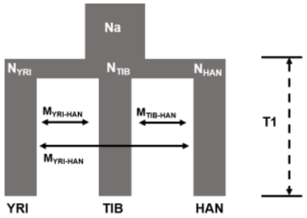 | All three populations differentiated at the same time, including Yoruba samples in Ibadan (YRI), Tibetans (TIB), Hans (HAN). | Na: 12,452<br>$N_{YRI}$: 11,187<br>$N_{TIB}$: 29,549<br>$N_{HAN}$: 31,190 | T1: 63,700 | $M_{YRI-TIB}$: 0.0879<br>$M_{TIB-HAN}$: 0.0018<br>$M_{YRI-HAN}$: 0.0882 | -453,165 | 906,345 |
| Model 2<br>(10 parameters) | 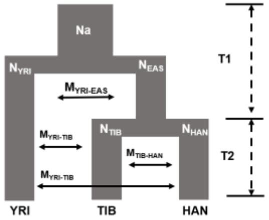 | YRI branched out first | Na: 12,495<br>$N_{YRI}$: 28,885<br>$N_{EAS}$: 8,906<br>$N_{TIB}$: 34,657<br>$N_{HAN}$: 43,446 | T1: 65,662<br>T2: 22,974 | $M_{YRI-EAS}$: 0.0166<br>$M_{YRI-TIB}$: 0.0983<br>$M_{TIB-HAN}$: 0.0072<br>$M_{YRI-HAN}$: 0.001 | -223,444 | 446,909 |
| Model 3<br>(17 parameters) | 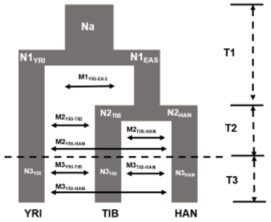 | YRI branched out first and population size changed | Na: 12,924<br>$N1_{YRI}$: 35,442<br>$N1_{EAS}$: 8,264<br>$N2_{TIB}$: 48,819<br>$N2_{HAN}$: 38,663<br>$N3_{YRI}$: 2,807<br>$N3_{TIB}$: 62,909<br>$N3_{HAN}$: 64,086 | T1: 62,144<br>T2: 22,414<br>T3: 1,274 | $M1_{YRI-EAS}$: 0.0063<br>$M2_{YRI-TIB}$: 0.0984<br>$M2_{TIB-HAN}$: 0.0013<br>$M2_{YRI-HAN}$: 0.001<br>$M3_{YRI-TIB}$: 0.0113<br>$M3_{TIB-HAN}$: 0.0035<br>$M3_{YRI-HAN}$: 0.0107 | -224,443 | 448,921 |

LL, log likelihood; AIC, Akaike information criterion.

**Table S7. Maximum likelihood parameter estimates and confidence intervals of best-fit demographic models.**

| Best-fit model for TIB-HANN-HANS | | | | Best-fit model for YRI-TIB-HAN | | | |
|---|---|---|---|---|---|---|---|
| **Parameters** | **Descriptions** | **Estimates** | **95% CIs** | **Parameters** | **Descriptions** | **Estimates** | **95% CIs** |
| $N_a$ | The size of the ancestral population to Tibetans and Hans | 10,206 | 9,987 – 10,425 | $N_a$ | The size of the ancestral population to Yoruba samples in Ibadan (YRI) and East Asians (EAS) | 12,495 | 11,931 – 13,059 |
| theta | Mutation rates per $4N_a$ generations | 98,384 | 96,273 – 100,494 | theta | Mutation rates per $4N_a$ generations | 130,826 | 124,917 – 136,734 |
| $N_{TIB}$ | The ancestral population size of Tibetans | 36,260 | 36,234 – 36,285 | $N_{YRI}$ | The ancestral population size of YRI | 28,885 | 28,863 – 28,906 |
| $N_{HAN}$ | The ancestral population size of Hans | 50,870 | 50,647 – 51,092 | $N_{EAS}$ | The ancestral population size of EAS | 8,906 | 8,855 – 9,435 |
| $N_{HANN}$ | The ancestral population size of Han in the North (HANN) | 50,667 | 50,485 – 50,848 | $N_{TIB}$ | The ancestral population size of Tibetans | 34657 | 34,605 – 34,776 |
| $N_{HANS}$ | The ancestral population size of Han in the South (HANS) | 1,632 | 1,626 – 1,637 | $N_{HAN}$ | The ancestral population size of Hans | 43,446 | 43,435 – 43,453 |
| T1 | The divergence time between Tibetans and Hans | 16,871 | 14,049 – 19,692 | T1 | The divergence time between YRI and EAS | 65,662 | 63,109 – 68,214 |
| T2 | The divergence time between HANN and HANS | 769 | 727 – 810 | T2 | The divergence time between Tibetans and Hans | 22,974 | 19,763 – 26,184 |
| $M_{TIB\text{-}HAN}$ | Migration rate between Tibetans and Hans per $2N_a$ generations | 0.0016 | 0 – 0.0032 | $M_{YRI\text{-}EAS}$ | Migration rate between YRI and EAS per $2N_a$ generations | 0.0166 | 0.0138 – 0.0193 |
| $M_{TIB\text{-}HANN}$ | Migration rate between Tibetans and HANN per $2N_a$ generations | 0.0087 | 0.0082 – 0.0091 | $M_{YRI\text{-}TIB}$ | Migration rate between YRI and Tibetans per $2N_a$ generations | 0.0983 | 0.0955 – 0.1010 |
| $M_{HANN\text{-}HANS}$ | Migration rate between HANN and HANS per $2N_a$ generations | 0.0537 | 0.0432 – 0.0641 | $M_{TIB\text{-}HAN}$ | Migration rate between Tibetans and Hans per $2N_a$ generations | 0.0072 | 0.0054 – 0.0089 |
| $M_{TIB\text{-}HANS}$ | Migration rate between Tibetans and HANS per $2N_a$ generations | 0.003 | 0 – 0.0175 | $M_{YRI\text{-}HAN}$ | Migration rate between YRI and Hans per $2N_a$ generations | 0.001 | 0.0002 – 0.0017 |

All scaled parameter estimates were transformed into real values using a mutation rate of $1.5 \times 10^{-8}$ per site per generation and a generation time of 29 years

**Table S8. Uniform priors for demographic parameters relevant to events prior to the anatomically modern humans.**

| **Parameters** | **Range** |
|---|---|
| The size of the ancestral population to chimpanzee and human ($N_{chimp-hominin}$) | unif(50,000, 100,000) |
| The contemporary population size of chimpanzee ($N_{chimp}$) | unif(10,000, 50,000) |
| The size of ancestral population to hominin ($N_{hominin}$) | unif(10,000, 20,000) |
| The ancestral population size of Denisovans, Neanderthals, and their common ancestors ($N_{ARC}$) | unif(2,000, 4,000) |
| The divergence time between chimpanzee and human ($T_{chimp-hominin}$) | unif(5,000,000, 10,000,000) |
| The divergence time between archaic hominins and humans ($T_{ARC-AMH}$) | unif(522,000, 634,000) |
| The divergence time between Denisovan and Neanderthal ($T_{DNS-NDL}$) | unif(392,000, 438,000) |
| The divergence time between Siberian and European Neanderthal ($T_{Altai-Vindija}$) | unif(130,000, 145,000) |

This table was extracted from Hsieh, et al. 2019. Time in years and population sizes are the number of individuals

**Table S16. Summary of structural variations (SVs) with signatures of both selection (p-value < 0.05 for iHS) and introgression (p-value < 0.05 for $f_d$-statistic).**

| No. | SV Location (GRCh37) | Type | Genotyped allele frequency | | | | | iHS (p-value) | $f_d$ (p-value) | |
|---|---|---|---|---|---|---|---|---|---|---|
| | | | Tibetans | Hans | Biaka | DNS | NDL | | DNS | NDL |
| 1 | chr9:81,725,770-81,726,090 | Deletion | 0.63 | 0.30 | 0.56 | 0.00 | 0.00 | 0.003 | 0.009 | 0.013 |
| 2 | chr7:1,185,068-1,187,658 | Deletion | 0.77 | 0.45 | 0.92 | **1.00** | **0.50** | 0.028 | 0.025 | 0.001 |
| 3 | chr18:45,099,900-45,100,212 | Deletion | 0.24 | 0.02 | 0.48 | 0.00 | 0.00 | 0.000 | NS | 0.049 |
| 4 | chr14:106,055,915-106,056,047 | Deletion | 0.10 | 0.36 | 0.00 | 0.00 | 0.00 | 0.047 | 0.019 | 0.044 |
| 5 | chr21:42,402,890-42,403,212 | Deletion | 0.70 | 0.39 | 1.00 | 0.00 | **0.25** | 0.045 | 0.046 | 0.001 |
| 6 | chr12:52,354,758-52,354,891 | Deletion | 0.60 | 0.29 | 0.04 | 0.00 | 0.00 | 0.049 | 0.019 | 0.039 |
| 7 | chr14:106,184,197-106,185,014 | Deletion | 0.38 | 0.13 | 0.15 | 0.00 | 0.00 | 0.047 | 0.006 | 0.044 |
| 8 | chr8:103,880,129-103,880,130 | Insertion | 0.34 | 0.10 | 0.08 | 0.00 | 0.00 | 0.034 | 0.026 | 0.025 |
| 9 | chr10:79,136,314-79,136,397 | Deletion | 0.61 | 0.33 | 0.25 | 0.00 | 0.00 | 0.042 | 0.005 | 0.047 |
| 10 | chr12:52,395,084-52,395,849 | Deletion | 0.61 | 0.36 | 0.21 | 0.00 | 0.00 | 0.049 | 0.019 | 0.039 |
| 11 | chr10:119,979,200-119,979,514 | Deletion | 0.22 | 0.05 | 0.63 | 0.00 | 0.00 | 0.005 | 0.002 | 0.014 |
| 12 | chr5:87,668,428-87,668,602 | Deletion | 0.09 | 0.28 | 0.46 | 0.00 | 0.00 | 0.033 | 0.043 | 0.036 |
| 13 | chr6:95,400,827-95,400,920 | Deletion | 0.93 | 0.75 | 0.19 | 0.00 | 0.00 | 0.006 | NS | 0.010 |
| 14 | chr7:158,809,629-158,809,695 | Deletion | 0.01 | 0.13 | 0.10 | 0.00 | 0.00 | 0.042 | NS | 0.037 |
| 15 | chr1:2,919,030-2,919,365 | Deletion | 0.06 | 0.23 | 0.00 | **0.50** | **0.50** | 0.026 | NS | 0.016 |

DNS, Denisovans; NDL, Neandertals; iHS, integrated haplotype homozygosity score; $f_d$, $f_d$-statistic which is designed to distinguish excess genetic drift from ancient introgression based on allele frequencies; NS, not significant.
